# Reproducible evaluation of transposable element detectors with McClintock 2 guides accurate inference of Ty insertion patterns in yeast

**DOI:** 10.1101/2023.02.13.528343

**Authors:** Jingxuan Chen, Preston J. Basting, Shunhua Han, David J. Garfinkel, Casey M. Bergman

## Abstract

**Background:** Many computational methods have been developed to detect non-reference transposable element (TE) insertions using short-read whole genome sequencing data. The diversity and complexity of such methods often present challenges to new users seeking to reproducibly install, execute or evaluate multiple TE insertion detectors.

**Results:** We previously developed the McClintock meta-pipeline to facilitate the installation, execution, and evaluation of six first-generation short-read TE detectors. Here, we report a completely re-implemented version of McClintock written in Python using Snakemake and Conda that improves its installation, error handling, speed, stability, and extensibility. McClintock 2 now includes 12 short-read TE detectors, auxiliary pre-processing and analysis modules, interactive HTML reports, and a simulation framework to reproducibly evaluate the accuracy of component TE detectors. When applied to the model microbial eukaryote *Saccharomyces cerevisiae*, we find substantial variation in the ability of McClintock 2 components to identify the precise locations of non-reference TE insertions, with RelocaTE2 showing the highest recall and precision in simulated data. We find that RelocaTE2, TEMP, TEMP2 and TEBreak provide a consistent and biologically meaningful view of non-reference TE insertions in a species-wide panel of ~1000 yeast genomes, as evaluated by coverage-based abundance estimates and expected patterns of tRNA promoter targeting. Finally, we show that best-in-class predictors for yeast have sufficient resolution to reveal a dyad pattern of integration in nucleosome-bound regions upstream of yeast tRNA genes for Ty1, Ty2, and Ty4, allowing us to extend knowledge about fine-scale target preferences first revealed experimentally for Ty1 to natural insertions and related copia-superfamily retrotransposons in yeast.

**Conclusion:** McClintock (https://github.com/bergmanlab/mcclintock/) provides a user-friendly pipeline for the identification of TEs in short-read WGS data using multiple TE detectors, which should benefit researchers studying TE insertion variation in a wide range of different organisms. Application of the improved McClintock system to simulated and empirical yeast genome data reveals best-in-class methods and novel biological insights for one of the most widely-studied model eukaryotes and provides a paradigm for evaluating and selecting non-reference TE detectors for other species.

## Background

Transposable elements (TEs) are mobile, repetitive DNA sequences that occupy large fractions of most eukaryotic genomes and can cause mutations of large effect [1]. Since their discovery, a variety of cytological, molecular and genomic techniques have been developed to identify TE insertions in genomes at varying levels of resolution [2, 3, 4, 5, 6, 7, 8]. In principle, the most comprehensive method for detecting new TE insertions in a genome is whole genome sequencing (WGS) followed by *de novo* genome assembly and systematic genome-wide annotation of TEs. However, the difficulty of accurately assembling repetitive DNA sequences using short reads [9, 10] and the challenges of generating high-quality long-read genome sequencing data hinder using assembly-based methods for genome-wide detection of TE insertions, especially for large population samples [11].

Currently, the most widely-used approach for detection of new TE insertions involves mapping unassembled short-read WGS data to a reference genome and inferring the location and family using split-read or read-pair information [8]. Over the last decade, many bioinformatic methods have been developed for the reference-based detection of TE insertions using short-read WGS data (https://tehub.org/en/resources/repeat_tools) [12]. Most of these methods were initially developed for a specific organism with its unique TE composition, and thus have different design goals and features. The general performance of short-read TE detectors is typically unknown because of the “self-assessment trap” and limited number of methods compared in primary studies [13]. A few independent performance evaluation studies have been conducted comparing subsets of short-read TE detectors, but these studies differ in their evaluation frameworks, selection of tools, and focal organisms [14, 15, 16]. Crucially, no study to date has provided a reproducible evaluation of multiple TE detectors, making it difficult to validate published results or extend findings beyond the context of the initial evaluation study. Despite these issues, one emerging theme from current evaluation studies is that considerable variation in performance exists among different short-read TE detectors, across organisms and evaluation frameworks [8, 14, 15, 16]. As such, there is not yet any clear basis to select the best short-read TE detector for a new organism, and thus researchers must base tool choice by extrapolation from performance on other taxa or employ multiple methods to ensure robust biological conclusions [17, 18, 19].

Previously, we developed a meta-pipeline called McClintock [15] to facilitate the installation and execution of six first-generation short-read TE detectors (ngs_te_mapper [20], PoPoolationTE [21], RelocaTE [22], RetroSeq [23], TE-locate [24], and TEMP [25]). The original McClintock system automated installation of these six “component” TE detection methods, provided a common interface to run all components, reduced the number of shared input files, and generated a standard set of output files [15]. Since its initial development, the McClintock system has been used to support detection of TE insertions and enable biological discoveries in a variety of organisms and biological contexts [15, 16, 17, 18, 19, 26, 27, 28, 29, 30, 31, 32, 33, 34, 35, 36, 37, 38, 39, 40, 41, 42, 43, 44] and to facilitate comparative evaluation of multiple TE detectors [15, 16, 45].

A long-term aim of the McClintock project was to develop a flexible framework to incorporate additional TE detectors as they were published, allowing researchers to run the most relevant set of methods and compare performance on their genomes of interest. While the original pipeline made it easier to install and run multiple TE detectors and compare their output, its initial design had several limitations that made it difficult to achieve this long-term aim. Most importantly, the original McClintock pipeline was implemented in Bash with minimal use of functions or classes to encapsulate, abstract, or modularize the code, and most variables were defined in the global scope. These shortcomings made it difficult to modify and expand the original codebase without compromising existing functionality, and thus hindered the addition of new TE detectors to the original framework.

The original McClintock system also had several limitations related to its installation. Component TE detectors were installed automatically by McClintock, however the software dependencies for each component had to be installed manually, which required substantial effort on the part of the user. Moreover, all components in the original McClintock system ran in a single computing environment, which necessitated a mutually-compatible set of software dependencies to be installed. This fragile configuration also caused compromises in the versions of software dependencies that were used and, in some cases, locked the original system to increasingly out-of-date versions of software dependencies.

Even after successful installation, the original McClintock system had a number of limitations regarding its usability. Notably, error handling was largely absent from the original codebase. Thus, when failures occurred during the execution of component TE detectors or their software dependencies, McClintock would continue to run, often producing a cascade of error messages from downstream processes that made it difficult for users to know how and why the pipeline failed. Similarly, the original pipeline had strict input file formatting requirements (e.g., requiring unzipped fastq files), but which would provide no warnings or messages if the input files did not comply to required specifications. Moreover, the original McClintock system hard-coded some input and post-processing parameters that were not easily modified by users but impacted the performance of component methods. The original McClintock pipeline also output TE insertion predictions only in Browser Extensible Data (BED) format (https://samtools.github.io/hts-specs/BEDv1.pdf), rather than the Variant Call Format (VCF) [46] that is standard for reporting genetic variation relative to a reference genome. Finally, the original pipeline also did not produce plots or tables to facilitate comparison of predictions across component methods.

To address these limitations, we completely re-implemented McClintock in Python leveraging Conda (https://github.com/conda/conda) and Snakemake [47] to improve the installation, error handling, speed, stability, and extensibility of the pipeline. We also incorporated six new TE detectors (ngs_te_mapper2 [35], PoPoolationTE2 [48], RelocaTE2 [49], TEBreak [50], TEFLoN [51], and TEMP2 [52]), doubling the total number of components available in the new McClintock system. In addition, the updated McClintock system now includes new modules to automate read trimming, provide estimates of TE family abundance from depth of coverage, and produce an integrated HTML report summarizing predictions made by all components. Importantly, McClintock now also provides a new reproducible simulation framework that allows users to evaluate the performance of component TE detectors in a flexible manner across different organismal contexts. In this report, we describe the structure and function of the updated McClintock system, highlight its improvements and new features, evaluate component TE detector performance using simulated and empirical yeast genome data, and use the updated McClintock system to provide new biological insights into the pattern of TE insertion in *S. cerevisiae*.

## Implementation

### Re-implementation of McClintock in Python using Conda and Snakemake

Here we describe the major features distinguishing the re-implemented McClintock 2 system from the original version reported in Nelson *et al.* [15] (Table 1). We initially sought to further develop the original Bash-based McClintock system reported in Nelson *et al.* [15] by improving its installation using the Conda package and environment management system (https://github.com/conda/conda). The use of Conda allows the creation of distinct environments (both for McClintock itself and for the component methods wrapped in McClintock) that can be reproducibly generated on different computing systems without the need for root privileges. This modification to the original McClintock system permitted external software dependencies to be automatically installed separately for each component, and each component to be executed in its own isolated environment. While solving some of the problems with the installation and versioning of external dependencies in the original system, this development work led to an endpoint because it did not solve most of the software engineering and usability limitations described above. Therefore, the final version of the Bash-based McClintock system extended to use Conda (referred to in this report as McClintock 1) was retired.

**Table 1.** Major features that distinguish the original McClintock version reported in Nelson et al. [15] and the McClintock 2 system reported in the current study.

|  | Nelson *et al.* [15] | McClintock 2 |
| --- | --- | --- |
| McClintock implementation | Bash | Python |
| Workflow management | N.A. | Snakemake |
| Dependency installation | Manual | Conda (automated) |
| # Component methods | 6 | 12 (with extensibility) |
| Read trimming and QC | N.A. | trimgalore module |
| TE copy number estimation | N.A. | coverage module |
| Output format | .BED | .VCF and .BED |
| Summary report | .CSV | .HTML (interactive) |
| Reproducible simulation system | N.A. | Built-in (with documentation) |

N.A. indicates that the feature is not available.

We next completely rewrote the McClintock system in Python 3 using Conda and Snakemake [47], with the initial aim of recapitulating the general functionality in McClintock 1. The new McClintock 2 codebase is heavily modularized and leverages the native error handling functions in Python. The Snakemake workflow manager was chosen to underpin the McClintock 2 system for multiple reasons. First, we could use the Snakemake system to simplify and fully automate installation of all components and their software dependencies (either from Conda channels like Bioconda [53] or directly from project repositories), and to automatically create independent Conda environments for each component. Second, the “meta-pipeline” architecture of McClintock is naturally suited to a rule-based workflow, with each component being encapsulated as an individual rule that is executed by Snakemake inside a specific Conda environment whose dependencies do not conflict with any other component/rule. Third, since the Snakemake engine determines the rules that are run to create the requested outputs, no additional coding logic is required in McClintock itself to manage the complexities of which components were run and in which order, or how files shared by multiple components are created and used (e.g., reference genomes, trimmed read files, BAM files, etc.). Importantly, this virtue of Snakemake applies regardless of how many components are executed at run time or incorporated into McClintock 2, permitting flexible execution of various components by users and easy addition of new components by developers. Finally, the Snakemake engine can run multiple independent rules simultaneously if enough central processing unit (CPU) cores are available, potentially allowing for multiple rules to be run in parallel on multi-core systems. In the re-implemented system, if *n*>1 cores are requested, McClintock 2 by default now allocates a maximum of *n*/2 (for an even number of cores) or (*n*–1)/2 (for an odd number of cores) to each rule which allows multiple rules to be executed in parallel. Optionally, users can specify the “–serial” flag which prevents multiple rules from being run in parallel and allows rules that use multiple cores to have access to maximal number of cores requested.

After integrating the original six components (ngs_te_mapper [20], PoPoolationTE [21], RelocaTE [22], RetroSeq [23], TE-locate [24], and TEMP [25]) into the McClintock 2 system, we cross-validated its accuracy versus the original McClintock 1 system using simulated genomes generated by our new reproducible simulation platform (see below and Supplemental Text). We then incorporated six more recently-developed TE detectors that fulfilled our original inclusion criteria [15] into the re-engineered McClintock system: ngs_te_mapper2 [35], PoPoolationTE2 [48], RelocaTE2 [49], TEBreak [50], TEFLoN [51], and TEMP2 [52].

In addition to providing a full re-implementation and additional TE detectors, McClintock 2 has new functionality not present in the original system (Table 1). First, McClintock 2 now has a preprocessing module to perform read trimming and quality control (QC) using TrimGalore (https://github.com/FelixKrueger/TrimGalore) and produce a read QC report using MultiQC [54]. If the read trimming option is set (either optionally in conjunction with user-specified components, or in a default run that executes all 12 components), then trimmed reads are used as input for other components in McClintock 2. Second, McClintock 2 has a new analysis module that can generate estimates of TE copy number using a “coverage module” that computes relative depth-of-coverage for each query TE sequence normalized by depth-of-coverage in non-repetitive regions of the genome. The coverage module also produces plots for each query that allow users to inspect variation in read depth across TE sequences, which may be caused by the existence of multiple TE sub-families that differ by structural variants (e.g., [17, 55]). Third, McClintock 2 reports non-reference TE insertion variants predicted by each component in VCF format [46] in addition to BED files containing standardized information about reference and non-reference TE insertions. Fourth, McClintock 2 is able to generate an interactive HTML report for each run that allows users to view, sort, and filter results for each component and TE family (Fig. 1). Finally, the McClintock 2 repository also provides code to automatically simulate TE insertions in any user-supplied genome and generate synthetic WGS datasets that can be used as input to evaluate component TE detector performance (detailed in the following section).

**Figure 1.**
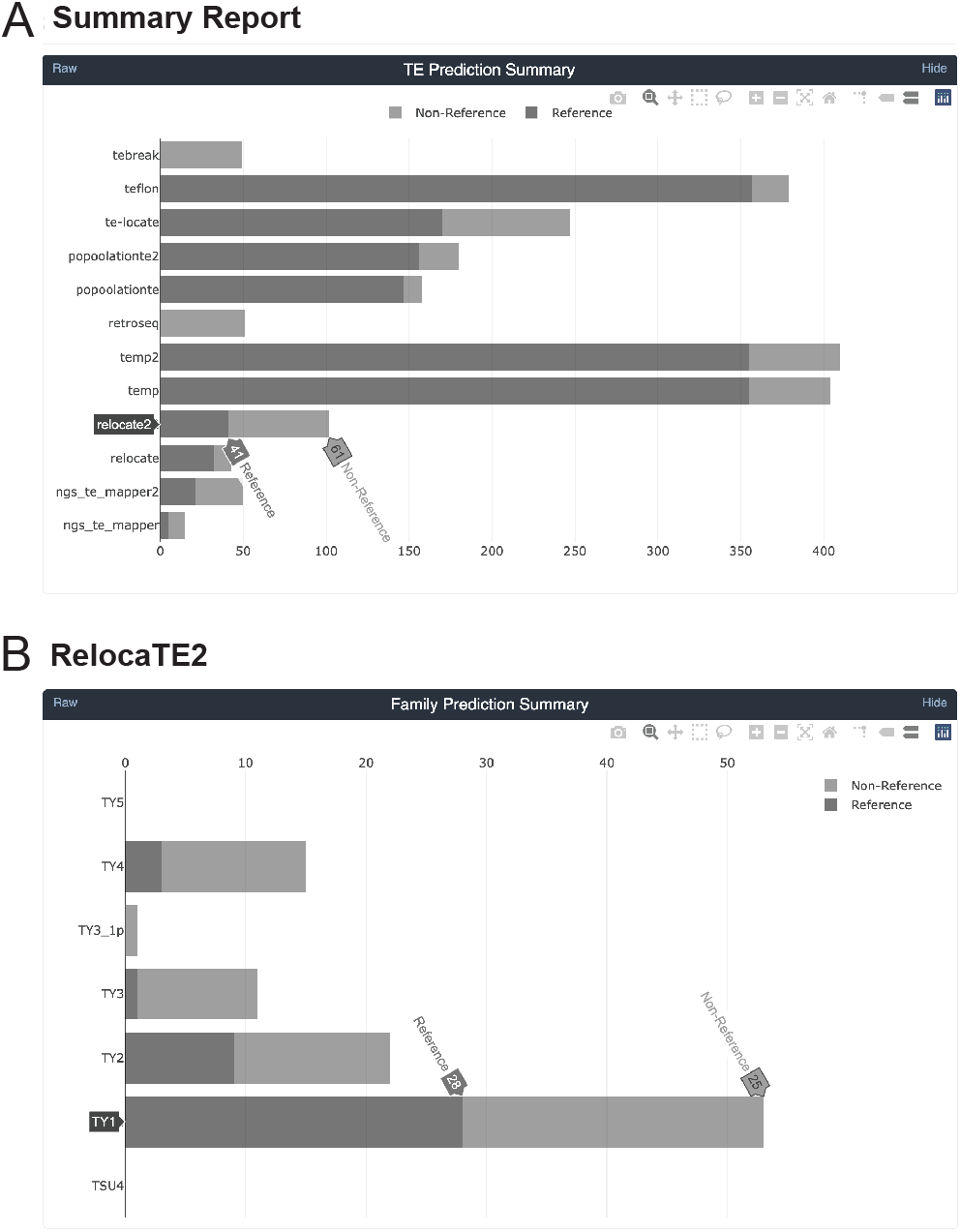
Sample screenshots from the new interactive HTML report in McClintock 2. The HTML report generates summary information for the McClintock run including interactive bar plots for: (A) the number of reference, non-reference, and total number of predictions made across all TE families by all 12 component methods; and (B) the number of reference, non-reference, and total number of predictions made for a specific component method (e.g., RelocaTE2). Barplots from the report shown were generated by a complete McClintock run (revision d2b819a18b2a549be483fdcc948e1346e589a4cb) applied to Illumina 101-bp paired-end sequences for *S. cerevisiae* strain YJM1460 (SRA: SRR800842), down-sampled to 50× fold-coverage.

### A reproducible simulation system for evaluating McClintock component performance

We previously developed a “single synthetic insertion” framework for evaluating the performance of McClintock components to detect non-reference TE insertions and used this framework to evaluate the performance of the six TE detectors in McClintock 1 [15]. This evaluation framework created a synthetic genome comprised of a single non-reference TE and its corresponding target site duplication (TSD) inserted into in an otherwise-unmodified reference genome, then simulated a corresponding paired-end WGS dataset that could be used as input for McClintock. The Bash code for this simulation framework was not released as part of the original McClintock 1 system, did not run inside a controlled environment, was hard-coded to simulate TE insertion preferences for only one species *(S. cerevisiae*), only evaluated only one WGS fold-coverage (100×), and did not generate standard performance metrics like precision and recall.

To overcome these limitations, we re-implemented a flexible and fully reproducible version of this single synthetic insertion simulation framework in Python 3 using Conda and Snakemake. The new simulation framework in McClintock 2 consists of three major parts that run under controlled Conda environments and allows customization for different organisms:

i. A python script that creates a configuration file with information about the simulation experiment to be run in JavaScript Object Notation (JSON) format. Required inputs to the configuration script include user-supplied files containing the reference genome, TE library, and permissible locations for TE insertions. The configuration script also allows users to modify the number of replicates in the experiment, properties of the synthetic genomes (e.g., TSD length and strand of the TE insertion), properties of the simulated WGS datasets (e.g., read length, single or paired end reads, insertion sizes for paired-end reads, read error rates, fold-coverage), which read simulator will be used to generate WGS datasets (ART [56] or wgsim [57]), and computing resources for each cluster job (e.g., number of threads, amount of memory).
ii. A primary Snakemake workflow that reads input parameters from the JSON file created by the configuration script, then submits and manages jobs on a high-performance computing (HPC) cluster. Currently our simulation framework is designed for use with cluster systems running the SLURM workload manager, but it can be adapted to other HPC systems using alternative Snakemake cluster profiles (https://github.com/Snakemake-Profiles/). Each submitted job runs a Python script that creates a new synthetic genome with one additional insertion, simulates a corresponding WGS dataset from this synthetic genome, then runs the McClintock pipeline on the simulated WGS dataset. Replicates for each combination of parameters are submitted as individual cluster jobs to maximize available cluster resources, and any potential failed jobs are automatically re-submitted by the workflow to minimize effort needed to monitor the simulation experiment.
iii. A secondary Snakemake workflow that is automatically executed after all cluster jobs submitted by the primary workflow are completed, which summarizes results and generates preliminary plots from all replicates of the overall simulation experiment. This workflow runs several scripts to calculate performance metrics (e.g., mean numbers of non-reference TE predictions, precision, recall) across replicates, generate precision/recall curves as a function of fold-coverage, and generate positional accuracy and predicted TSD length plots for each fold-coverage level. Performance metrics are calculated as follows. Nonreference TE predictions with the identical family, chromosome, and strand, with start and end locations falling within an N bp window from the synthetic insertion (N=0, 5, 100, 300, or 500) were labeled as “within-N” predictions. Within-0 bp predictions have the identical start and end coordinates as the synthetic insertion in that simulated sample and are considered “exact” predictions. The mean number of reference and non-reference TE predictions (overall, and at various window sizes) per synthetic genome were then calculated across all replicates. To calculate precision and recall, non-reference TE predictions within-N bp from the insertion site that match the TE family and strand of the synthetic insertion were regarded as true positives (TP), while all other predictions were regarded as false positives (FP). If more than one nonreference prediction for the correct TE family and strand was made within-N bp from the synthetic insertion, only one was considered a TP and all others were considered FPs. If a replicate had zero non-reference TE predictions within-N bp from the synthetic insertion, it was regarded as a false-negative (FN). For each N bp window size, recall was calculated as TP/(TP+FN) and precision was calculated as TP/(TP+FP).

## Materials and Methods

### Evaluation of McClintock component performance on simulated data

We conducted a series of simulation experiments to validate aspects of the McClintock re-implementation, demonstrate the utility of our new reproducible simulation framework, and evaluate the ability of McClintock components to predict nonreference TE insertions in *S. cerevisiae*. The different simulation experiments varied in the version of McClintock used, which component methods were run, their placement of synthetic insertions, the read simulator used to generate WGS datasets, and performance metrics used (See Supplemental Text for details). In all cases, synthetic genomes were created with the new reproducible simulation framework available in McClintock 2 (see Implementation above), the UCSC sacCer2 version of the *S. cerevisiae* S288c reference genome (to allow cross-validation with results in Nelson *et al.* [15]), canonical sequences for *S. cerevisiae* Ty elements from [58] (https://github.com/bergmanlab/mcclintock/blob/master/test/sac_cer_TE_seqs.fasta), and 5-bp TSDs for all Ty families [15, 59, 60, 61, 62]. Likewise, all McClintock jobs that were run on simulated data used the UCSC sacCer2 version of the *S. cerevisiae* S288c reference genome with reference TE annotations, taxonomy files, and canonical sequences for *S. cerevisiae* Ty sequences from [58] (provided in https://github.com/bergmanlab/mcclintock/blob/master/test/) as input. Quantitative results from all simulations can be found in Additional File 2 (average numbers of TEs) and Additional File 3 (recall and precision).

### Evaluation of McClintock run times on empirical data

To test whether Snakemake’s ability to run multiple rules simultaneously could improve run-time performance in the new McClintock system, we applied the original McClintock 1 (revision 714fe6d6aa04a7ccdb0d26718e08960783a6229a) and the new McClintock 2 (revision f766e73a8d04efdbd57664bc5195abc022806674) systems to a 101-bp paired-end Illumina HiSeq 2000 dataset for *S. cerevisiae* strain YJM1460 [63] that was down-sampled to 50× and 100× coverage using seqtk (v1.3) [64]. To compare run times for McClintock 1 and McClintock 2, we invoked both versions of the system only with the original six components present in McClintock 1 (“-m ngs_te_mapper,relocate,temp,retroseq,popoolationte,te-locate”). To evaluate the impact on run time caused by addition of the six new components in McClintock 2, we also invoked McClintock 2 using all 12 components (“-m ngs_te_mapper,ngs_te_mapper2,relocate, relocate2,temp,temp2,retroseq,popoolationte, popoolationte2,te-locate,teflon,tebreak”). In both cases, we varied the number of cores requested (1, 5, 10, 15, 20, 25, 30) but kept the amount of RAM constant (20 Gb). For all runs, we used unzipped, untrimmed fastq reads as input (as required by McClintock 1), the UCSC sacCer2 version of the *S. cerevisiae* S288c reference genome, reference TE annotations, taxonomy files, and canonical TE sequences of each *S. cerevisiae* Ty family from [58]. Trials were run serially on the same cluster node (an AMD EPYC processor with a total of 32 cores and 256GB RAM, operating system Linux 3.10.0-1160.36.2.el7.x86_64). The average and standard deviation of CPU efficiency and run times was calculated from five replicates of each setting (defined as the combination of number of cores and McClintock run options). All possible settings were run as a batch for each replicate to control for differences in cluster node performance over the duration of the experiment.

### Analysis of TE insertions in a diverse panel of wild and domesticated yeast strains

To demonstrate the utility of the McClintock 2 system for large-scale mining of empirical short-read WGS data, we applied our pipeline to Illumina paired-end datasets for 1,011 *S. cerevisiae* isolates [63, 65] sequenced to variable depths (from 50× to 900× fold-coverage). Illumina WGS data were downloaded from NCBI Sequence Read Archive and converted to fastq format using SRA toolkit (v2.10.8) [66]. To reduce the influence of variable coverage and computing resources, if the coverage of the original dataset was greater than 50×, we down-sampled it to 50× fold-coverage using seqtk (v1.3) [64]. The full McClintock 2 pipeline using all 12 components plus the coverage and trimgalore module (“-m ngs_te_mapper,ngs_te_mapper2,relocate, relocate2,temp,temp2,retroseq,popoolationte,popoolationte2,te-locate,teflon,tebreak, coverage,trimgalore”) (revision 7aa529881e72299af928a1a38cf809fdbd8e8bb3) was applied to this yeast resequencing dataset using the UCSC sacCer2 version of the *S. cerevisiae* S288c reference genome, reference TE annotations, taxonomy files, canonical sequences of each *S. cerevisiae* Ty family from [58].

We identified non-reference TE insertions in the vicinity of tRNA genes using BEDtools window (v2.30.0,“-u -sw -l 1000 -r 500”) [67]. BEDtools closest (“-D b”) was used to calculate the genomic distance between each non-reference TE insertion and its closest tRNA transcription start site (TSS). To analyze the relationship between the distribution of non-reference TE insertions and nucleosome occupancy upstream of tRNA genes, we mapped micrococcal nuclease digestion with deep sequencing (MNase-seq) data from [68] to the sacCer2 version of the *S. cerevisiae* S288c reference genome using Bowtie (v1.2.3 [69]) and generated genomewide nucleosome occupancy profiles using NUCwave [70]. Whole genome nucleosome occupancy profiles in .wig format were converted to .bw format using UCSC wig-ToBigWig (v377), and then Bwtool (v20170428, “bwtool aggregate 2000:500”) [71] was used to calculate nucleosome occupancy in the region 2 kb upstream to 500 bp downstream of all tRNA TSSs.

### Data analysis and visualization

Data were analyzed and visualized in R (v3.6.3) using the ggplot2 (v3.3.3) [72] and GGally (v2.1.1, https://ggobi.github.io/ggally/) packages. Kernel density plots showing distributions of non-redundant Ty insertion sites around tRNA genes were created with the ggplot2 function “geom_density” with Gaussian kernel and 0.4× default bandwidth.

## Results and Discussion

### Validation of McClintock 2 implementation and simulation system

Here we present a Python-based re-implemention and extension of the McClintock TE detection meta-pipeline, as well as a reproducible simulation system to test the performance of component TE detectors integrated into McClintock. The major features that distinguish the new McClintock 2 system from the original version reported in Nelson *et al.* [15] are described in the Implementation section above and are summarized in Table 1. To validate the McClintock 2 implementation and simulation system, we conducted a series of *in silico* experiments testing the ability of the six original components to detect a single synthetic non-reference TE insertion introduced into an otherwise unmodified *S. cerevisiae* reference genome (see Supplemental Text for details). Contrasts between different simulations allowed us to test whether various improvements in the new McClintock 2 system could yield results that replicate those previously published in Nelson *et al.* [15] including: (i) the Python-based implementation of the simulation system, (ii) the Python-based implementation of the meta-pipeline, and (iii) the random insertion model in the single insertion simulation framework. Results from these simulations (Fig. S1) allowed us to validate that the McClintock 2 re-implementation yields results that broadly replicate those previously published in Nelson *et al.* [15].

### Parallelization by Snakemake in McClintock 2 improves multi-core job run times

McClintock 2 uses Snakemake as a workflow manager, which allows parallelization of multiple rules when enough CPU cores are available. To investigate potential improvements in computational efficiency due to Snakemake job management in McClintock 2, we executed McClintock 1 and McClintock 2 (using either the six original or all 12 components) with a variable number of cores under controlled computing environments. For this analysis, we used a 101-bp paired-end Illumina dataset from the *S. cerevisiae* domesticated palm-wine strain Y12 (YJM1460; SRA: SRR800842) downsampled to either 50 × or 100× (see Materials and Methods for details). In general, we observe a decreasing trend in CPU efficiency for all McClintock run configurations as the number of CPU cores increases (Fig. 2A). However, when more than one core is used, we found that McClintock 2 has better CPU efficiency than McClintock 1 for the six original components, and that CPU efficiency is highest for all numbers of cores used when executing all 12 McClintock 2 components. Improved CPU efficiency for McClintock 2 is observed at both 50× and 100× coverage.

**Figure 2.**
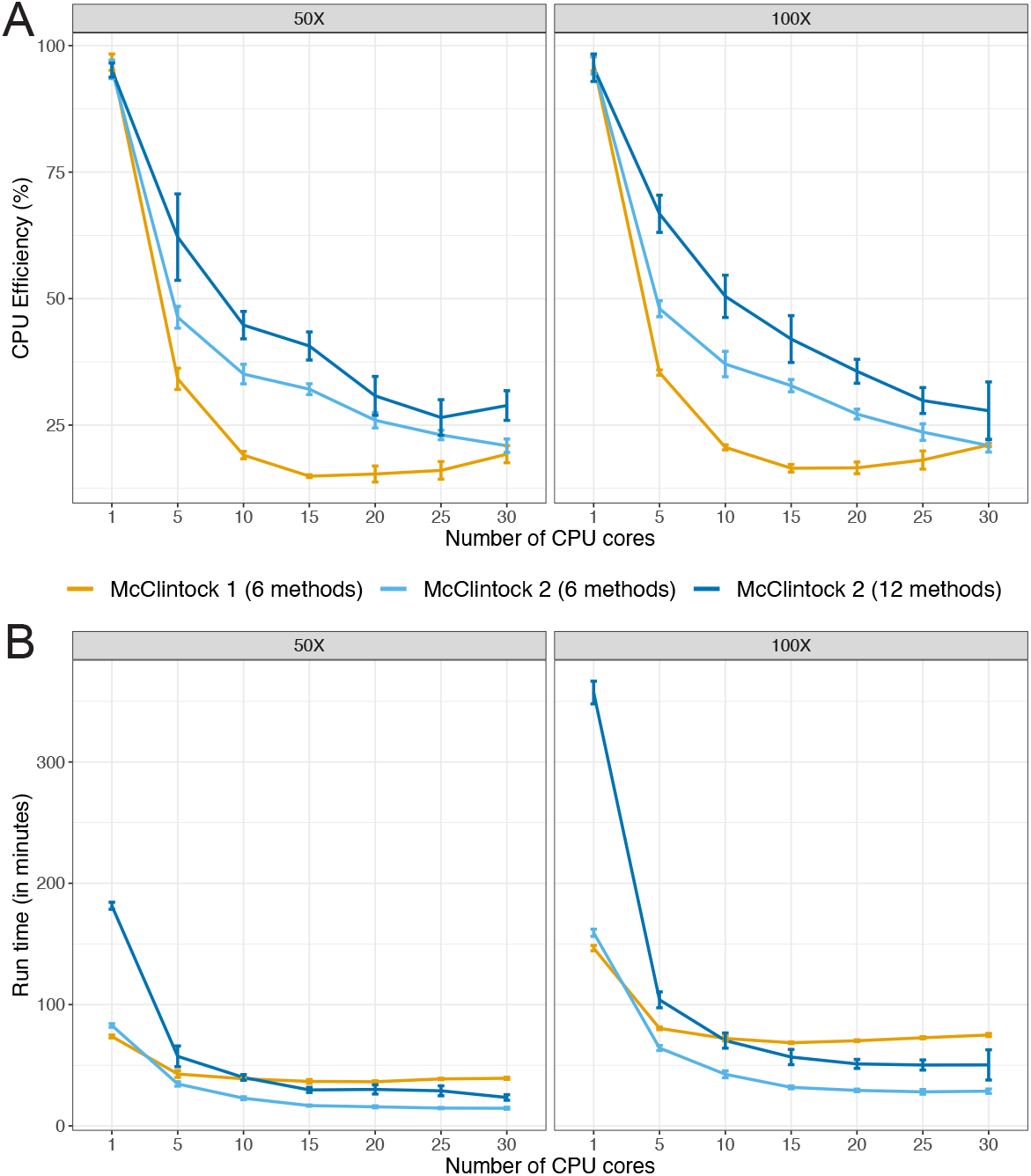
McClintock 2 re-implementation improves CPU efficiency and run time on multi-core architectures. Shown are average (A) CPU efficiency and (B) run times across 5 replicates of McClintock 1 (six component methods, orange line) or McClintock 2 (same six component methods as for McClintock 1, light blue line; all 12 component methods in McClintock 2, dark blue line) applied to 50× and 100× Illumina 101-bp paired-end sample for S. *cerevisiae* strain YJM1460 (SRA: SRR800842). Error bars indicate standard deviations across replicates. To allow compatibility with McClintock 1, all runs were performed on unzipped, untrimmed fastq files and thus run times do not include these processes.

When comparing run times for the six original components (Fig. 2B), we found that McClintock 2 was slightly slower (~10%) than McClintock 1 when using only a single core, however McClintock 2 finished increasingly faster when five or more cores were available. Longer run times were observed for McClintock 2 when executing all 12 components relative to only the six original components, regardless of the number of cores used. When running either the original six or all 12 components, McClintock 2 run times continued to decrease up to 15 allocated cores. Interestingly, when 15 or more cores are allocated, executing McClintock 2 with all 12 components is faster than running McClintock 1 using only six components, demonstrating clear speed improvements in the re-engineered version of the McClintock system relative to the original. Together, these results indicate that parallelization afforded by using Snakemake in McClintock 2 results in better utilization of computing resources in multi-core computing systems relative to McClintock 1.

### Evaluation of McClintock 2 component TE detectors on simulated yeast data

McClintock 2 now includes six additional component methods for detecting nonreference TE insertions using short-read WGS data that are not available in McClintock 1: ngs_te_mapper2 [35], PoPoolationTE2 [48], RelocaTE2 [49], TEBreak [50], TEFLoN [51], and TEMP2 [52]). Four of the new components (ngs_te_mapper2, PoPoolationTE2, RelocaTE2, TEMP2) are “second-generation” versions of tools previously incorporated in McClintock 1, and the other two (TEBreak and TEFLoN) represent new TE detection strategies not represented in McClintock 1.

To evaluate the relative performance of these 12 TE detection strategies, we used the new reproducible single insertion simulation approach available in McClintock 2. The evaluation of McClintock 2 component methods performed here differs from the approach originally used in Nelson *et al.* [15] in several ways (see Supplemental Text for details). First, we used a more biologically realistic model to randomly select the family and location of simulated non-reference TE insertions over a range of genomic positions. Second, we investigated the effects of component performance over a range of fold-coverages (3×, 6×, 12×, 25×, 50×, 100×), which provides practical guidance for users to optimize data generation. Third, the summary statistics used in Nelson *et al.* [15] are not standard evaluation metrics, and therefore difficult to compare with other evaluation studies (e.g., [14, 16]). Thus, here we used the standard evaluation metrics of recall and precision derived from the number of truepositive, false-positive, and false-negative predictions at various window sizes (see Implementation for details). Finally, here we evaluated performance using simulations that model yeast TE target preferences (Simulation 3) as in Nelson *et al.* [15], as well as random insertion into non-repetitive DNA (Simulation 4). Simulation 4 helps us interpret how insertion into repetitive DNA in the reference genome influences component performance in yeast and gives insight into potential component performance in other organisms that don’t have strong TE targeting preferences.

As shown previously for the original six components in McClintock 1 [15], different methods in McClintock 2 vary in their ability to accurately predict the position of synthetic TE insertions in simulated data (Figs. S2, S3, S4, S5). Several components (i.e., ngs_te_mapper, ngs_te_mapper2, RelocaTE, RelocaTE2, TEFLoN and TEBreak) make most predictions very close the expected location of the synthetic insertion, whereas other components (i.e., PoPoolationTE, PoPoolationTE2, and TE-locate) predict insertions over a wider distance around the synthetic insertion (~500bp). Even for components that attempt to identify insertion breakpoints at nucleotide resolution using split-read information, only ngs_te_mapper and RelocaTE2 have predictions with the expected 5-bp TSD length for all Ty families (Figs. S6, S7). This method-dependent variation in positional accuracy makes it difficult to fairly compare performance of different components at a single window size. Therefore, we calculated performance metrics over a range of window sizes for overlaps between predicted and simulated insertions (exact, within-5, within-100, within-300, and within-500 bp). Inspection of the complete set of results for all window sizes (Figs. S8, S9, S10, S11) revealed that two window sizes (exact and within-100 bp) could illustrate the majority of key features in McClintock 2 component performance across the insertion models and range of fold-coverages investigated here.

To guide selection of TE detectors in the model species *S. cerevisiae*, we first analyzed the performance of McClintock 2 components using data from Simulation 3 where TEs were inserted in their biologically realistic locations in yeast upstream of tRNA genes [73, 74, 75, 76, 77] (Fig. 3; purple lines). Three general trends can be observed in the recall and precision curves for McClintock 2 components for nonreference TEs in tRNA promoter regions. First, component performance is higher when allowing a more relaxed window size (within-100 bp) to classify TP predictions (dashed lines) than when requiring TP predictions to be exact (solid lines), consistent with most methods making many predictions that are not precise to exact nucleotide coordinates. Only ngs_te_mapper makes all its predictions exactly, and thus performance curves are overlayed for both window sizes shown for this method. Second, component performance typically increases with sequencing depth and then plateaus at a method-specific coverage. Some exceptions to this trend are observed for PoPoolationTE recall and RelocaTE and TEFLoN precision, which show decreasing performance with increasing coverage. Performance curves for most components suggest that 25× is the most cost-effective depth of sequencing coverage, and 50× is sufficient to optimize performance for most component methods. Third, second generation versions of TE detectors – apart from PoPoolationTE2 – typically show improved performance relative to their first generation counterparts, suggesting that gains in TE detector performance can be made by continued method development. Overall, results from Simulation 3 suggest that RelocaTE2 has remarkably high recall (~92%) and precision (~98%) to predict non-reference TE insertions in tRNA promoter regions of the yeast genome (at 50× coverage and ?5 bp window size). The recall and precision of TEMP, TEMP2, RetroSeq and TEBreak to detect TE insertions in yeast promoter regions are all both greater than 75% when allowing non-exact predictions in WGS datasets with greater than 50× coverage.

**Figure 3.**
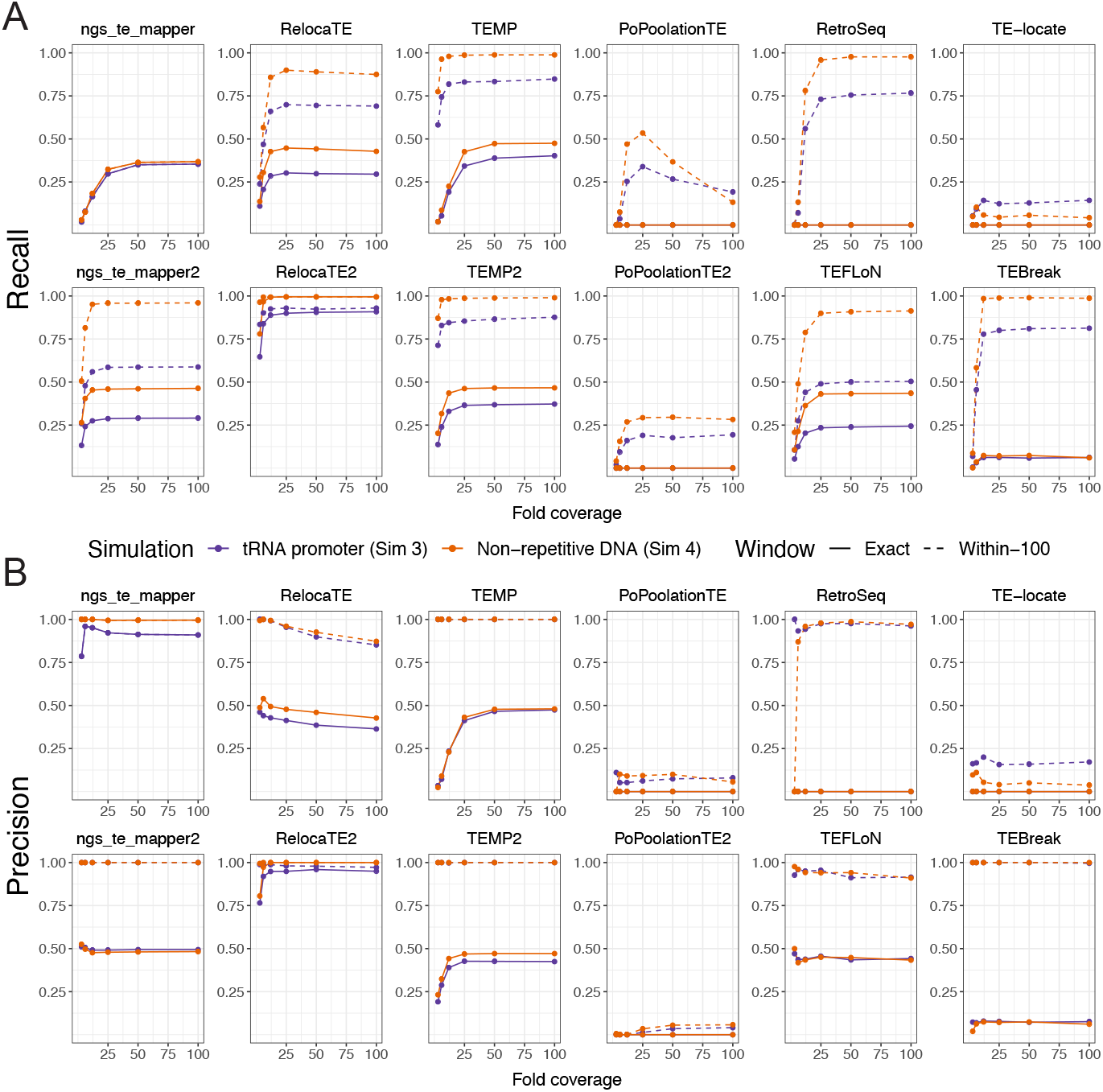
Performance of McClintock 2 component methods in simulated yeast WGS data. Shown are the (A) recall and (B) precision across different fold-coverage for individual compnent methods to detect single synthetic insertions in an otherwise unmodified S. *cerevisiae* reference genome. Purple lines (Simulation 3) model the biologically realistic insertion preferences of yeast TEs, with synthetic Ty insertions created upstream of tRNA genes in regions that often have fragments of prior TE insertions in the reference genome. Orange lines (Simulation 4) model random insertions in non-repetitive regions, which allows insight into the effects of insertion within repetitive DNA and component’s performance for organisms without strong TE targeting preferences. Points indicate tested fold-coverage configurations, i.e, 3×, 6×, 12×, 25×, 50× and 100 ×. Solid lines represent performance estimates for non-reference TE predictions made at the exact site of the synthetic insertion. Dashed lines represent performance estimates for non-reference TE predictions made within 100 bp surrounding the synthetic insertion site. The six original component methods in McClintock 1 are on the top row of each panel, and the six new methods in McClintock 2 are on the second row of each panel.

While biologically relevant for *S. cerevisiae*, the tRNA promoter insertion model used in Simulation 3 potentially underestimates component performance since synthetic insertions are often placed into fragments of TE sequences that occur upstream of tRNA genes in the reference genome [58, 62]. To gain insight into other organismal contexts and understand how insertion into reference TE sequences may impact McClintock 2 component performance in yeast, we also investigated a model of random insertion into unique regions of the *S. cerevisiae* genome (Simulation 4). Recall and precision curves for the random insertion model (Fig. 3; orange lines) show the same general trends but have consistently higher performance relative to results for the same component in the tRNA promoter insertion model (Fig. 3; purple lines). General performance improvements in Simulation 4 relative to Simulation 3 indicate that prediction of non-reference TE insertions into locally TE-rich regions (like tRNA promoters in yeast) is more challenging for most component methods than for those in non-repetitive DNA. Consistent with results from the tRNA promoter model, RelocaTE2 has the highest recall and precision of any component in McClintock 2, with essentially perfect performance in non-repetitive regions of the *S. cerevisiae* genome (~100%) even at exact base-pair resolution. At the within-100 bp window size, TEMP, TEMP2, RetroSeq and TEBreak also perform very well at identifying non-reference insertions in unique regions. We note that two additional methods – ngs_te_mapper2 and TEFLoN– show dramatically better recall for non-exact predictions in non-repetitive regions relative to tRNA promoter regions, putting these methods in a similar performance class to TEMP, TEMP2, RetroSeq and TEBreak for insertions in non-repetitive DNA.

In summary, results from our reproducible simulation experiments show that RelocaTE2 is the best-performing component method currently in the McClintock 2 meta-pipeline to detect non-reference TE insertions in *S. cerevisiae* under both the biologically relevant tRNA promoter and non-repetitive insertion models. If exact base-pair positional accuracy is not required, TEMP, TEMP2, RetroSeq and TEBreak also exhibit high performance to detect TE insertions in *S. cerevisiae* tRNA promoter regions at 50× coverage. Furthermore, our results suggest that at 50× coverage TEMP, TEMP2, RetroSeq, TEBreak, ngs_te_mapper2, and TEFLoN may have high performance to detect TE insertions in non-repetitive regions of other genomes. Finally, our results indicate that users should provide McClintock 2 with WGS datasets of at least 25× fold-coverage to generate the best performance from most component methods, but that coverage higher than 50× may not lead to further performance benefits.

### Best-in-class component methods reveal consistent insights into patterns of TE insertion in yeast

Our reproducible simulation system allowed us to validate the McClintock 2 re-implementation and evaluate component methods under ideal conditions, where only one TE insertion per strain is to be detected and all sequencing data is generated in an identical way. However, analysis of large-scale empirical re-sequencing data is expected to present additional challenges to TE detectors, such as more complex insertion patterns in a given strain (e.g., multiple insertions, tandem elements) and variability in WGS dataset composition across strains (e.g., fold-coverage or median insert size). To assess component method performance under a more realistic population genomic scenario, we applied the complete McClintock 2 system (all 12 components plus the coverage module) to empirical WGS datasets from a world-wide sample of 1,011 S. *cerevisiae* strains isolated from diverse environments and geographical locations [63, 65]. Since these WGS datasets varied significantly in their fold-coverage (50× to 900×), we down-sampled all WGS datasets to 50× prior to running McClintock 2 based on simulation results above that indicated 50 × coverage is sufficient to maximize component method performance (Fig. 3). Our goals for this analysis were to evaluate whether McClintock 2 component methods: (i) predict consistent numbers of non-reference TE insertions per strain (overall and by Ty family), and (ii) have sufficient positional accuracy to identify expected patterns of targeting in yeast tRNA promoter regions. This analysis also allowed us to generate the first species-wide Ty insertion variant call sets for *S. cerevisiae* (Additional File 4).

To evaluate consistency among component methods on empirical WGS data, we first quantified the distribution of non-reference TE predictions made by each method across 1,011 yeast strains (Fig. 4A). We observe substantial variation in median values of non-reference Ty insertions across all 12 methods, as well as in the correlation among methods in the number of predicted non-reference Ty insertions (Fig. S12). Interestingly, median values of ~50 non-reference Ty insertions per strain are consistently predicted by the five component methods (RelocaTE2, TEMP, TEMP2, RetroSeq and TEBreak) (Fig. 4A; bold outlines) that showed the best performance (when considering non-exact predictions) in simulated data under the biologically relevant tRNA promoter model (Fig. 3A; purple dotted lines). These five components also have the highest pairwise correlation of non-reference Ty counts per strain across methods (Fig. S12). We note that the original six components do not show the general pattern of ~50 non-reference Ty insertions per strain (Fig 4A; top row) and have some of the lowest correlation of non-reference Ty counts across methods (Fig. S12), underscoring the value of incorporating additional recently-developed methods into McClintock 2 to reveal consistent biological patterns that could not be revealed by the McClintock 1 system. Since McClintock 2 component methods do not distinguish full-length elements from solo LTRs, the emergent estimate of ~50 non-reference Ty insertions per yeast strain is likely to represent a combination of both structural types. Indeed, applying the McClintock 2 coverage module to internal coding regions of all Ty families, we estimate that the number of full-length elements per strain is ~20 (Fig. S13), suggesting that many non-reference Ty insertions detected by McClintock 2 components in this population sample are polymorphic solo LTRs [58, 78].

**Figure 4.**
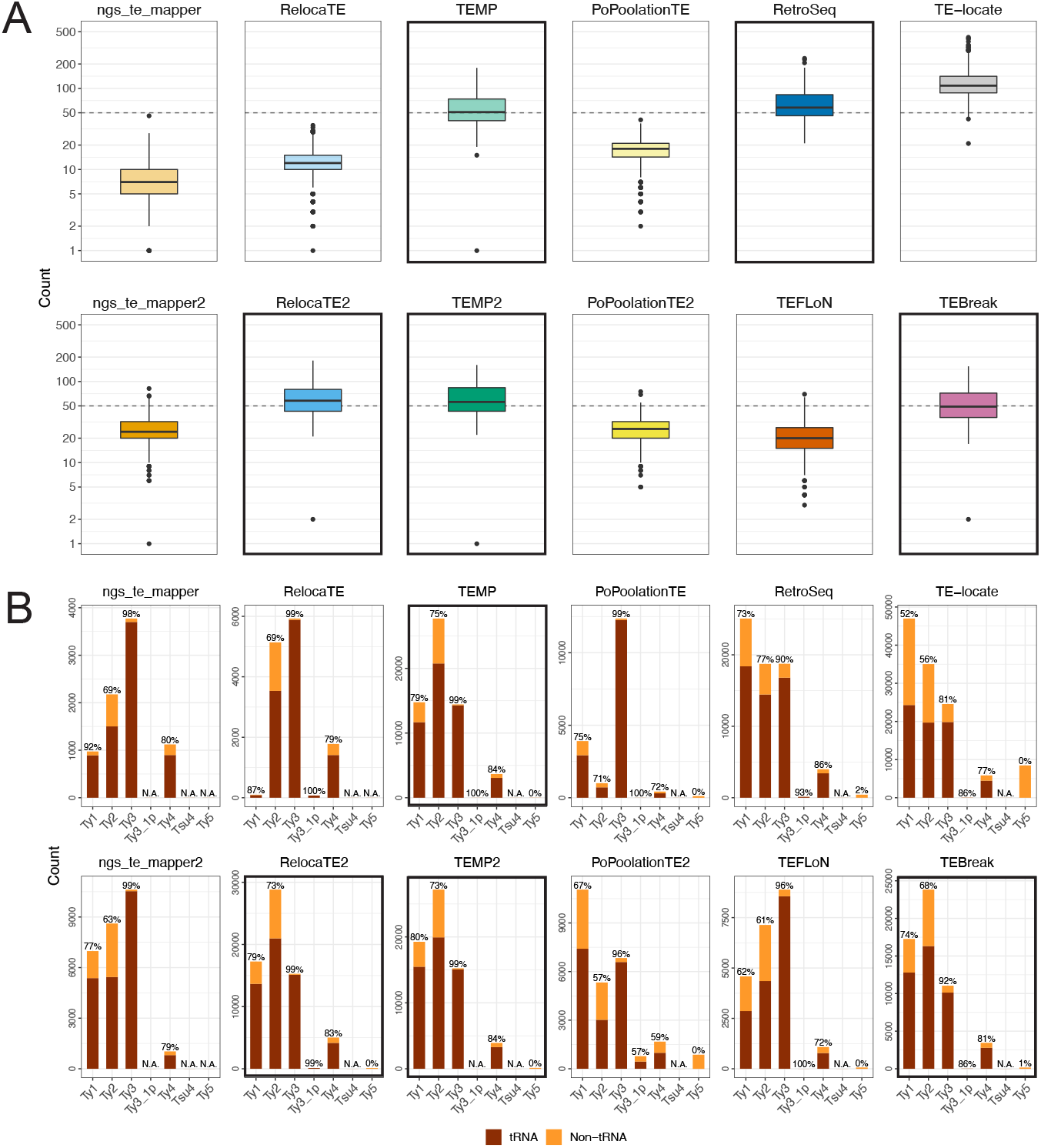
Numbers of Ty elements predicted by McClintock 2 components in a world-wide sample of yeast strains. (A) Numbers of non-reference TE predictions per strain (summed over all Ty families) and (B) numbers of non-reference TE predictions across Ty families (summed over all strains) in 1,011 S. *cerevisiae* WGS samples [63, 65], down-sampled to 50× fold-coverage. In panel (A), lines inside boxes indicate median values, colored boxes show interquartile ranges (IQR), whiskers show values 1.5× IQR of the upper or lower quartiles, and the dots indicate outliers that beyond 1.5×IQR. Components with bold outlines in panel (A) have have median values of ~50 non-reference Ty insertions per strain (dashed lines), as well as recall and precision both >75% in tRNA promoter insertion simulations when allowing non-exact predictions in WGS datasets with >50× coverage (see Fig. 3). We note that the y-axis is on a log_10_ scale, and that 16 zero-count data points and one extreme TE-locate data point (count=749) is removed to aid with visualization. In panel (B) total numbers of non-reference TE predictions are partitioned as “tRNA” (dark red) if they are located between 1000 bp upstream and 500 bp downstream of tRNA genes, or “non-tRNA” (orange) if outside these windows. Note that the y-scale varies for each component method. The percentage of near tRNA gene predictions is annotated at the top of each bar. “N.A.” means no such Ty family was found using that component. Components with bold outlines in panel (B) predict consistent relative TE family abundance and also have properties of components with bold outlines in panel (A), and thus we designate them as “best-in-class” methods for predicting non-reference TE insertions in *S. cerevisiae*.

Next, we assessed the consistency among component methods to classify the TE family of non-reference predictions across all 1,011 isolates (Fig. 4B). As for estimates of non-reference Ty abundance per strain, we observe variable patterns in relative Ty family abundance across component methods, with most methods being able to differentiate Ty families that are known to be active (Ty1, Ty2, Ty3 and Ty4) from those that are inactive or low-abundance (Ty3_1p, TSU4, Ty5). As observed for numbers of non-reference TEs per strain above, we note that no clear pattern of Ty family abundance can be discerned across the original six components (Fig. 4B; top row). However, a consistent pattern of relative Ty family abundance is observed for four of the five component methods (RelocaTE2, TEMP, TEMP2 and TEBreak) that show both consistent non-reference TE abundance per strain and high performance in simulated data. These four methods all indicate that Ty2 has the highest overall number of non-reference Ty insertions in this sample of *S. cerevisiae* strains. While Ty1 is often cited as the most abundant Ty family in *S. cerevisiae* because of its high copy number in the reference strain S288c [62, 58], the finding that Ty2 has the highest number of non-reference insertions in this diverse worldwide sample of *S. cerevisiae* strains is supported by orthogonal copynumber estimates from the McClintock 2 coverage module (Fig. S14) and a similar depth-based approach used in recent independent study [79]. In contrast, the fifth component method that shows consistent Ty abundance per strain and high performance in simulated data – RetroSeq – predicts more Ty1 and Ty2 insertions in empirical yeast data, which we infer to be misidentification because of the similarity in LTR sequences for these two Ty families [80, 62].

Previously [15], we used the well-established pattern that Ty1, Ty2, Ty3 and Ty4 non-randomly target promoters of genes transcribe by RNA polymerase III such as tRNAs (reviewed in [81]) to validate non-reference TE predictions made by McClintock 1 component methods in small sample of 93 *S. cerevisiae* genomes [15, 63]. Here, we confirm that the majority of non-reference predictions made by all component methods in McClintock 2 for these four Ty families are in the expected vicinity of tRNA genes in the species-wide *S. cerevisiae* resequencing dataset analyzed here (Fig. 4B) [63, 65]. Moreover, the four methods (RelocaTE2, TEMP, TEMP2 and TEBreak) that consistently predict similar numbers and families of non-reference Ty insertions in empirical yeast genome data, also predict similar proportions of insertions in tRNA genes. Combined with their high performance in simulated data, we conclude that RelocaTE2, TEMP, TEMP2 and TEBreak are the “best-in-class” methods for predicting Ty insertions in *S. cerevisiae* WGS resequencing data.

Finally, we sought to test whether predictions in species-wide *S. cerevisiae* WGS data made by the best-in-class components in McClintock 2 have sufficient positional accuracy to recapitulate the association between Ty1 insertion profiles and nucleosome occupancy in tRNA promoters observed for experimentally induced insertions [74, 76, 82]. Recent work by Hays *et al.* [40] using WGS data from experimentally-evolved genomes has shown that Ty insertions predicted by RelocaTE2, aggregated across all Ty families, exhibit periodicity upstream of tRNA genes in *S. cerevisiae* [40]. However, their analysis does not partition insertions to individual Ty families, show a direct correlation with nucleosome profiles, or demonstrate that this signal is not an artifact of a single TE detection system. As shown in Fig. 5, naturally-occurring Ty1 insertions predicted in species-wide *S. cerevisiae* WGS data by the four “best-in-class” component methods show a clear profile of dyad peaks on the first nucleosome-bound regions upstream of tRNA TSS, with weaker dyad signals on the second and third upstream nucleosome-bound regions. Intriguingly, Ty2 and Ty4 insertions predicted by all four best-in-class methods (and by RetroSeq, ngs_te_mapper2 and TEFLoN; Figs. S15 and S16) also have dyad peaks on the first nucleosome-bound region upstream of *S. cerevisiae* tRNA genes. Additionally, our results are support the observation that the domain of Ty1 integrase shown to be responsible for tRNA targeting is conserved in Ty2 and Ty4 [83]. In contrast, naturally-occurring Ty3 insertions show no correlation with nucleosomal profiles and are restricted to ~15 bp upstream of tRNA TSSs as seen for experimentally-induced insertions [77]. TEMP and TEMP2 predict another peak of insertions for Ty1, Ty2, and Ty4 at a similar position to the Ty3 peak, which is not observed in the insertion profiles for RelocaTE2 and TEBreak or in experimental data for Ty1 [74, 76, 82]. We interpret this additional peak at ~15 bp upstream of tRNA TSSs in the TEMP and TEMP2 methods as an artifact that is possibly caused by misassignment of some Ty3 insertions to other tRNA-targeting Ty families. Nevertheless, these results from natural genomes confirm that the insertion profiles for Ty1 and Ty3 upstream of tRNA genes are not artifacts of experimental induction, and suggest that the molecular mechanisms responsible for the targeting of Ty1 to nucleosomes are conserved in the related *copia*-superfamily retrotransposons Ty2 and Ty4.

**Figure 5.**
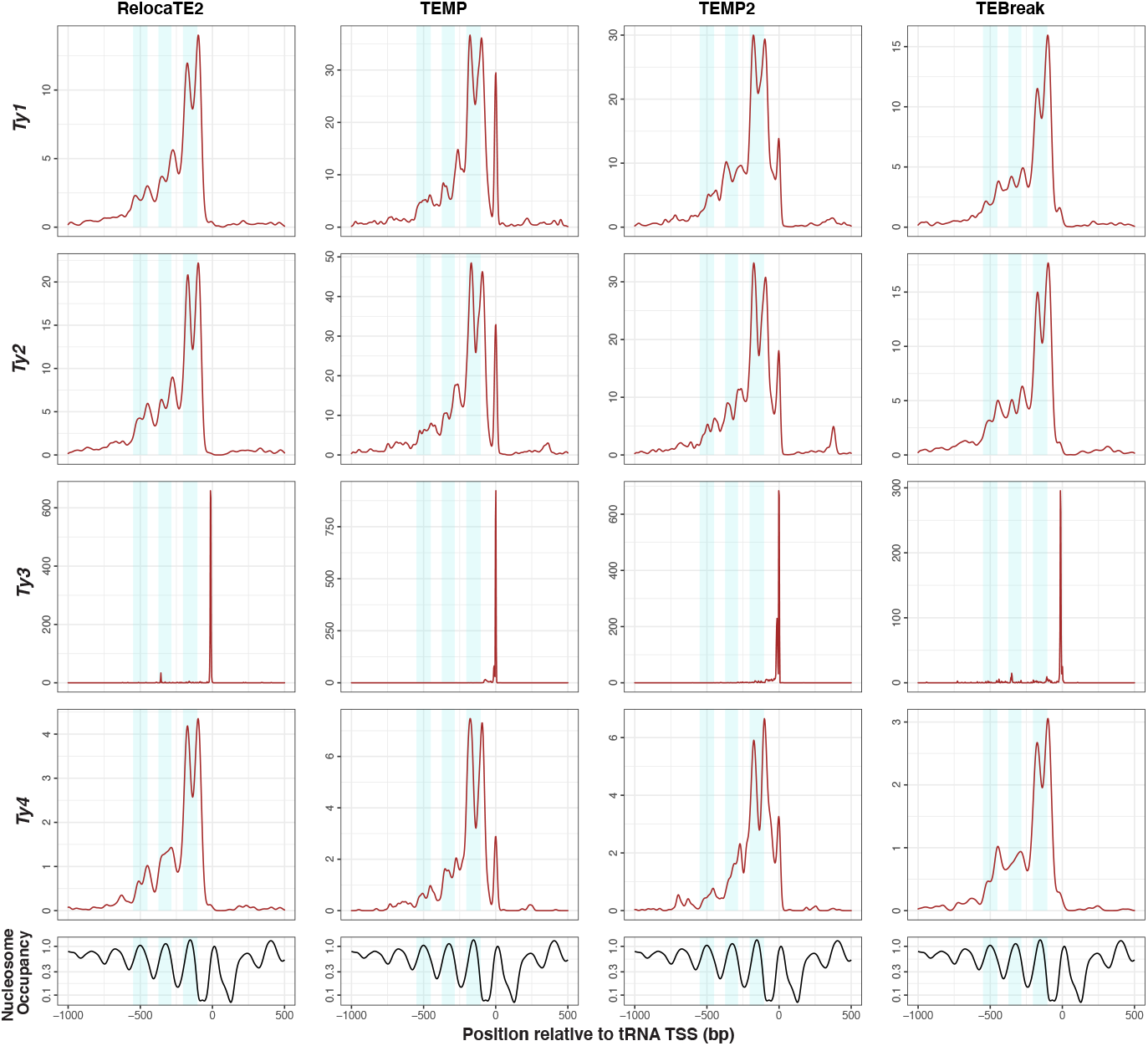
copia-superfamily retrotransposons show a dyad pattern of insertion in nucleosome-bound regions upstream of yeast tRNA genes. The top four rows show density profiles of non-redundant insertion sites for non-reference Ty predictions made by best-in-class McClintock 2 components (RelocaTE2, TEMP, TEMP2 and TEBreak) in tRNA promoter regions in a panel of 1,011 S. *cerevisiae* WGS samples [63, 65], down-sampled to 50× fold-coverage. Only the four Ty familes (Ty1, Ty2, Ty3 and Ty4) that are know to non-randomly target tRNA genes are included in this analysis. The bottom row shows nucleosome occupancy inferred using MNase-seq data from [68]. Light blue shaded areas indicate 100-bp regions surrounding peaks of nucleosome occupancy.

## Conclusions

In this report we present a re-implemented version of the McClintock TE detection system with improved installation, error handling, speed, stability, and extensibility. Relative to the original system [15], McClintock 2 includes six new short-read TE detectors, pre-processing and analysis modules, interactive HTML reports, and a reproducible simulation system that we used here to cross-validate the re-implemented system and evaluate component TE detectors. We acknowledge that not all available short-read TE detectors are currently incorporated into the McClintock 2 metapipeline, however use of Conda and Snakemake in the re-implemented meta-pipeline puts us in a better position to integrate additional or improved components in the future. We are also considering modifications to allow installation of TE detectors from user-downloaded executables or source code to be able to integrate methods like MELT [84] that have high performance but have restrictive licenses which prevent automatic installation. Our reproducible simulation framework now allows evaluation of McClintock 2 component performance in a wide range of organismal contexts besides *S. cerevisiae*, including humans and other model organisms. In the future, we plan to modify the simulation framework to allow multiple insertions in a single synthetic genome to reduce the number of replicates needed to analyze larger genomes that have longer run times. We also plan to implement a complementary evaluation system that uses TE annotations in long-read assemblies as gold standards to empirically benchmark TE detectors using short-read WGS data from the same strain [16, 52].

Based on performance evaluations in simulated WGS data, we conclude that RelocaTE2 is the best McClintock 2 component to detect non-reference TE insertions in *S. cerevisiae* at base-pair accuracy, which supports use of this method in recent yeast experimental evolution studies [85, 40]. However, we caution against the general conclusion that RelocaTE2 is the best McClintock 2 component for other species, since this method can have long run times that precludes its use in larger genomes [16, 52]. By combining results from simulated data with consistency analyses in empirical WGS data, we also identify TEMP, TEMP2 and TEBreak as additional “best-in-class” methods for predicting Ty insertions in *S. cerevisiae* WGS resequencing data with 50× fold-coverage or higher, albeit with lower positional accuracy than RelocaTE2. Results from the four best-in-class McClintock 2 components for *S. cerevisiae* provide additional support to the emerging view that Ty2 is the most abundant TE in this species [79], although we acknowledge that non-independence of strains and sampling bias in the strain panel analyzed [63, 65] could influence the conclusion about the relative abundance of Ty families in *S. cerevisiae*. Finally, we show that best-in-class McClintock 2 components for *S. cerevisiae* generate reasonable non-reference TE predictions with sufficient resolution to reveal a dyad pattern of integration in nucleosome-bound regions upstream of yeast tRNA genes for Ty1, Ty2, and Ty4. This finding allows knowledge about fine-scale target preferences revealed in experimentally induced Ty1 insertions [74, 76, 82] to be extended to natural Ty1 insertions and related *copia*-superfamily retrotransposons in yeast. Together with the new bioinformatics resources provided in McClintock 2, our work provides novel biological insights about TEs in one of the best-studied model systems and a paradigm for selecting optimal non-reference TE detectors in a diversity of organisms.

## Acknowledgements

We thank members of the Bergman and Garfinkel Labs for helpful discussions throughout the project; the BioConda community for contributing software recipes for software dependencies used in this project; and the Georgia Advanced Computing Resource Center for computing resources and HPC support.

## Abbreviations

BED: browser extensible data
CPU: central processing unit
FN: false negative
FP: false positive
HPC: high-performance computing
IQR: inter-quaritle range
JSON: JavaScript object notation
QC: quality control
TE: transposable element
TP: true positive
TSD: target site duplication
TSS: transcription start site
VCF: variant call format
WGS: whole genome sequencing

## Availability of data and materials

Output from McClintock runs on simulated and empirical yeast data can be found in Additional Files 2-4.

## Authors’ contributions

P.J.B. developed the McClintock 2 meta-pipeline with contributions from J.C., S.H. and C.M.B. J.C. and P.J.B. developed the reproducible simulation framework. J.C. conducted the analysis of simulated and empirical yeast genomes. J.C., P.J.B., and C.M.B analyzed the data. C.M.B. conceived of and managed the project. J.C., P.J.B. and C.M.B. drafted the manuscript with contributions from S.H. and D.J.G. All authors revised and approved the final manuscript.

## Additional Files

Additional File 1 — Supplemental Text, Tables and Figures. Combined PDF of all Supplemental Text, Figures and Tables.

Additional File 2 — Average numbers of TEs predicted by McClintock component methods in simulated data. CSV file with average numbers of reference and non-reference TEs predicted by McClintock 1 or 2 component methods in replicate simulations of yeast genomes with a single additional synthetic insertion, overall and at varying window size distances relative to the synthetic insertion.

Additional File 3 — Recall and precision for McClintock 2 component methods in simulated data. CSV file with numbers of true positive, false positive, and false negative non-reference TEs predicted by McClintock component methods (and corresponding recall and precision values) in replicate simulations of yeast genomes with a single additional synthetic insertion, at varying window size distances relative to the synthetic insertion.

Additional File 4 — Non-reference TEs predicted by McClintock component methods in 1,011 *S. cerevisiae* isolates. Multi-sample VCF files for non-reference TEs predicted by McClintock 2 component methods in 1,011 S. *cerevisiae* isolates.

