## Additional File 1 for "Reproducible evaluation of transposable element detectors with McClintock 2 guides accurate inference of Ty insertion patterns in yeast"

Supplementary material for

### Reproducible evaluation of short-read transposable element detectors and species-wide data mining of insertion patterns in yeast.

Jingxuan Chen[*,1], Preston J. Basting[*,1], Shunhua Han[*], David J. Garfinkel[†], & Casey M. Bergman[*,‡,2]

[*] Institute of Bioinformatics, University of Georgia, 120 E. Green St., Athens, GA, USA.
[†] Department of Biochemistry and Molecular Biology, University of Georgia, 120 E. Green St., Athens, GA, USA.
[‡] Department of Genetics, University of Georgia, 120 E. Green St., Athens, GA, USA.

[1] These authors contributed equally to this work.

### Supplementary Text

#### McClintock 2 cross-validation and performance evaluation simulation methods

We conducted four simulation experiments to validate aspects of the McClintock re-implementation, demonstrate the utility of our new reproducible simulation framework, and evaluate the ability of McClintock components to predict non-reference TE insertions in *S. cerevisiae*. In four simulations, synthetic genomes were created with the new reproducible simulation framework available in McClintock 2, the UCSC sacCer2 version of the *S. cerevisiae* S288c reference genome (to allow cross-validation with results in Nelson *et al.* [1]), canonical sequences for *S. cerevisiae* Ty elements from [2] (`https://github.com/bergmanlab/mcclintock/blob/master/test/sac_cer_TE_seqs.fasta`), and 5-bp TSDs for all Ty families [1, 3, 4, 5, 6]. Likewise, all McClintock jobs run on simulated data used the UCSC sacCer2 version of the *S. cerevisiae* S288c reference genome with reference TE annotations, taxonomy files, and canonical sequences for *S. cerevisiae* Ty sequences from [2] (provided in `https://github.com/bergmanlab/mcclintock/blob/master/test/`) as input. Details of the four simulation experiments are as follows:

Simulation 1 aimed replicate previous results reported in the original McClintock paper [1] using the new simulation framework. Simulation parameters were the same as in Nelson *et al.* [1] and are based on the well-established preferences of Ty elements to insert upstream of genes transcribed by RNA Pol III such as tRNAs (reviewed in [7]). To create synthetic genomes, a single Ty element was sequentially chosen from the list of Ty families that are active in S288c (Ty1, Ty2, Ty3, and Ty4) and inserted upstream one of the 299 tRNAs in reference genome. Potential insertion sites for Ty1, Ty2 and Ty4 were set to be 195-200 bp upstream of tRNA genes, and potential insertion sites for Ty3 were set to be 12-17 bp upstream of tRNA genes. In total, 598 synthetic genomes were generated: 299 genomes each having one insertion upstream of one tRNA gene in the forward strand, and 299 genomes each having one insertion upstream one tRNA gene in the reverse strand. As in Nelson *et al.* [1], the wgsim read simulator (v1, build h7132678_5) [8] was then used to create WGS paired-end datasets with a coverage of 100×, read lengths of 101 bp, median insert sizes of 300 bp and a base error rate of 1%. Performance for the six original components was evaluated using McClintock 1 (revision ad0c6cbaf9941328ac77953ec538c9604a141d3b).

Simulation 2 aimed to cross-validate the accuracy of the McClintock re-implementation using predictions from McClintock 2 for the six original components available in McClintock 1. Construction of synthetic genomes and simulation of WGS read datasets was the same as in Simulation 1. Performance for the six original components was evaluated using predictions from McClintock 2 (revision 7aa529881e72299af928a1a38cf809fdbd8e8bb3).

Simulation 3 aimed to evaluate the ability of all 12 components in McClintock 2 to detect TE insertions placed in more biologically-realistic locations in the *S. cerevisiae* genome over a range of WGS coverages. Simulation 3 used modified Ty targeting preferences and a different read simulator than in Simulations 1 and 2. First, rather than fixing the location of insertions for Ty1, Ty2, and Ty4 to be at one position (195-200 bp upstream of tRNA genes), we allowed synthetic insertions for these families to be placed in a broader region (50-800 bp upstream of tRNA genes) that better reflects patterns observed for both natural and induced insertions [1, 9, 10, 11]. As above, potential insertion sites for Ty3 were set to be 12-17 bp upstream of tRNA genes as is observed for both natural and induced insertions [1, 5, 12, 13]. Second, the tRNA gene and Ty family was chosen randomly rather than sequentially from the list of tRNA genes and Ty families. Also, because of incompatibilities between the format of fastq sequences generated by wgsim [8] and one of the new components in McClintock 2 (TEbreak), we used the ART read simulator (v2016.06.05 build h874f42a_8) [14] for Simulation 3. Simulated WGS paired end datasets were generated by ART with a built-in profile of Illumina HiSeq 2500 instrument, read lengths of 101 bp, median insert sizes of 300 bp, and a standard deviation of simulated DNA fragment sizes of 10 bp (options: -ss HS25 -sam -p -s 10 -l 101 -m 300). For each synthetic genome, six sets of simulated reads were produced at different fold-coverages (3×, 6×, 12×, 25×, 50× and 100×). We also rounded the number of replicates per strand to 300 for each fold-coverage. Performance for all 12 components was evaluated using predictions from McClintock 2 (revision 7aa529881e72299af928a1a38cf809fdbd8e8bb3).

Simulation 4 aimed to evaluate the performance of all 12 components in McClintock 2 to detect TE insertions in random, unique genomic regions over a range of WGS coverages. Because Ty targeting of RNA Pol III transcribed regions has caused a historical accumulation of Ty fragments upstream of tRNA genes [6, 2], many synthetic non-reference Ty insertions in Simulations 1-3 are placed into fragments of pre-existing Ty sequences found in the reference genome. Simulation 4 therefore allows us to gain insight into component method performance for TE families that do not target repetitive DNA in the reference genome (unlike Ty elements in *S. cerevisiae*), and by contrasting with results from Simulation 3 to understand the effects that insertion into repetitive DNA has on TE detector performance in yeast. Unique regions were defined as the complement of regions annotated as Ty elements in the UCSC sacCer2 version of the *S. cerevisiae* S288c reference genome [2]. Aside from differences in the location of potential insertion sites, construction of synthetic genomes and simulation of WGS read datasets was the same as in Simulation 3. Performance for all 12 components was evaluated using predictions from McClintock 2 (revision 7aa529881e72299af928a1a38cf809fdbd8e8bb3).

Quantitative results from all four simulations can be found in Additional Files 2 and 3.

#### Cross-validation of the McClintock 2 meta-pipeline and simulation system

To validate the new McClintock implementation and simulation system, we used results from Simulations 1-3 (see above). To be able to compare with previous results, we used the same approach as in Nelson *et al.* [1] to summarize the six original component methods' performance across simulated samples. Namely, we calculated the average number of non-reference TE predictions made by each component overall and at different levels of positional accuracy determined by varying window sizes (within 0, 5, 100, 300 or 500 bp of the synthetic insertion). Because differences in component performance on the positive and negative strands were minimal (Additional File 2), here we simplified reporting of summary metrics relative to Nelson *et al.* [1] by averaging results from forward and reverse strands across simulations (Fig S1). The expected value of non-reference TE insertions per simulation is one, with values less than one representing a tendency for a TE detector to make false negative predictions and values greater than one representing a tendency for false positive predictions.

We initially asked whether we could replicate performance results for the original six McClintock components reported in Nelson *et al.* [1] using the new reproducible simulation system in McClintock 2. This simulation (Simulation 1) used identical TE insertion settings but a more recent version of McClintock 1 and different simulation implementation relative to Nelson *et al.* [1] (Table S1). As shown in Fig. S1, results from Simulation 1 generated very similar numbers ($\pm 0.09$) of average non-reference TE predictions as in Nelson *et al.* [1], with the exception of the total number of non-reference TE predictions made by RelocaTE being substantially lower ($\sim$25%) in the current study. This difference could arise from many sources (including different git revisions for McClintock 1, component method dependencies, computing environments, simulation code, or run-to-run variation in component methods; Table S1) and underscores the difficulty in fully reproducing previous TE detector benchmarking studies. Nevertheless, the general consistency between results reported previously by Nelson *et al.* [1] and those generated independently here (Simulation 1) indicates that our new Python-based reproducible simulation system yields similar results as the old Bash-based simulation system, and that performance estimates of the original six component methods generated by McClintock 1 are broadly replicable.

Next, we sought to test whether the new Python-based McClintock 2 meta-pipeline generates similar performance results as the original Bash-based McClintock system. Because of the differences observed between Nelson *et al.* [1] and Simulation 1 (Fig. S1), we chose to use the reproducible results for McClintock 1 generated here (Simulation 1) as a baseline to compare with the McClintock 2 meta-pipeline. Also, using Simulation 1 as a baseline fully controls for differences between the new Python-based simulation system and the Bash-based system used in Nelson *et al.* [1], and therefore allows us to isloate any potential performance differences to the McClintock meta-pipeline itself. As shown in Fig. S1, no major differences in performance of the six orignal component methods ($\pm 0.065$) are observed for the Python-based (Simulation 2) and Bash-based (Simulation 1) implementations of the McClintock meta-pipeline. The largest difference observed is for RetroSeq, where this component exhibits improved performance in McClintock 2. Overall, this analysis demonstrates that performance estimates generated by the Python-based McClintock 2 are the same or better as those from the Bash-based McClintock 1 and indicates that no major differences should be experienced by users migrating from McClintock 1 to McClintock 2.

The approach used to simulate non-reference TE insertions in Nelson *et al.* [1] had a number of hard-coded constraints tailored for the yeast genome that made it unsuitable for application to other organismal contexts. Namely, the TE family and location of simulated insertions were generated sequentially from a candidate set of TE families and tRNA gene promoters, and the insertion positions were at fixed distances upstream of tRNA genes. To allow more general modeling of insertion preferences, we designed the new McClintock 2 simulation system to generate an instance of a randomly selected TE family to be randomly inserted into a set of arbitrary genomic locations specified in a configurable JSON file. By modifying parameters in the JSON file, the new simulation system is flexible enough to permit implementation of the Nelson *et al.* [1] approach as well as more biologically-realistic models of TE insertion, such as Ty insertion over a range of positions upstream of tRNA genes (see Materials and Methods for details). We tested if this new biologically-realistic insertion framework (Simulation 3) gave similar results at the same coverage (100$\times$) as the original sequential insertion framework with fixed Ty insertion locations (Simulation 2). As shown in Fig. S1, both the random (Simulation 3) and sequential (Simulation 2) insertion frameworks give similar results ($\pm 0.17$) with the exception of PoPoolationTE. We speculate that the unusually high number of total non-reference predictions made by PoPoolationTE in Simulation 3 may be caused by misclassification of reference TE insertions as non-reference TE insertions [1].

In summary, we can conclude that key improvements in McClintock 2 including (i) the Python-based implementation of the simulation system (Simulation 1 vs. Nelson *et al.* [1]), (ii) the Python-based implementation of the meta-pipeline (Simulation 2 vs. Simulation 1), and (ii) the random insertion model in the single insertion simulation framework (Simulation 3 vs. Simulation 2) together generate results that broadly replicate those published previously in Nelson *et al.* [1].

### Supplementary Tables

Table S1: Distinguishing features of simulation experiments performed in Nelson *et al.* [1] and this study.

| | Nelson *et al.* [1] | Simulation 1 | Simulation 2 | Simulation 3 | Simulation 4 |
|---|---|---|---|---|---|
| McClintock implementation | Bash | Bash | Python | Python | Python |
| McClintock git revision | e945d20 | ad0c6cb | 7aa5298 | 7aa5298 | 7aa5298 |
| Simulation implementation | Bash | Python | Python | Python | Python |
| Simulation git revision | N.A. | 7aa5298 | 7aa5298 | 7aa5298 | 7aa5298 |
| TE insertion targets | tRNA promoter | tRNA promoter | tRNA promoter | tRNA promoter | Non-repetitive DNA |
| Insertion position | Fixed distance | Fixed distance | Fixed distance | Specified range | Specified range |
| Candidate selection | Serial | Serial | Serial | Random | Random |
| Read simulator | wgsim | wgsim | wgsim | ART | ART |
| Simulated fold-coverage | 100× | 100× | 100× | 3× to 100× | 3× to 100× |
| Replicates per setting | 299 | 299 | 299 | 300 | 300 |

### Supplementary Figures

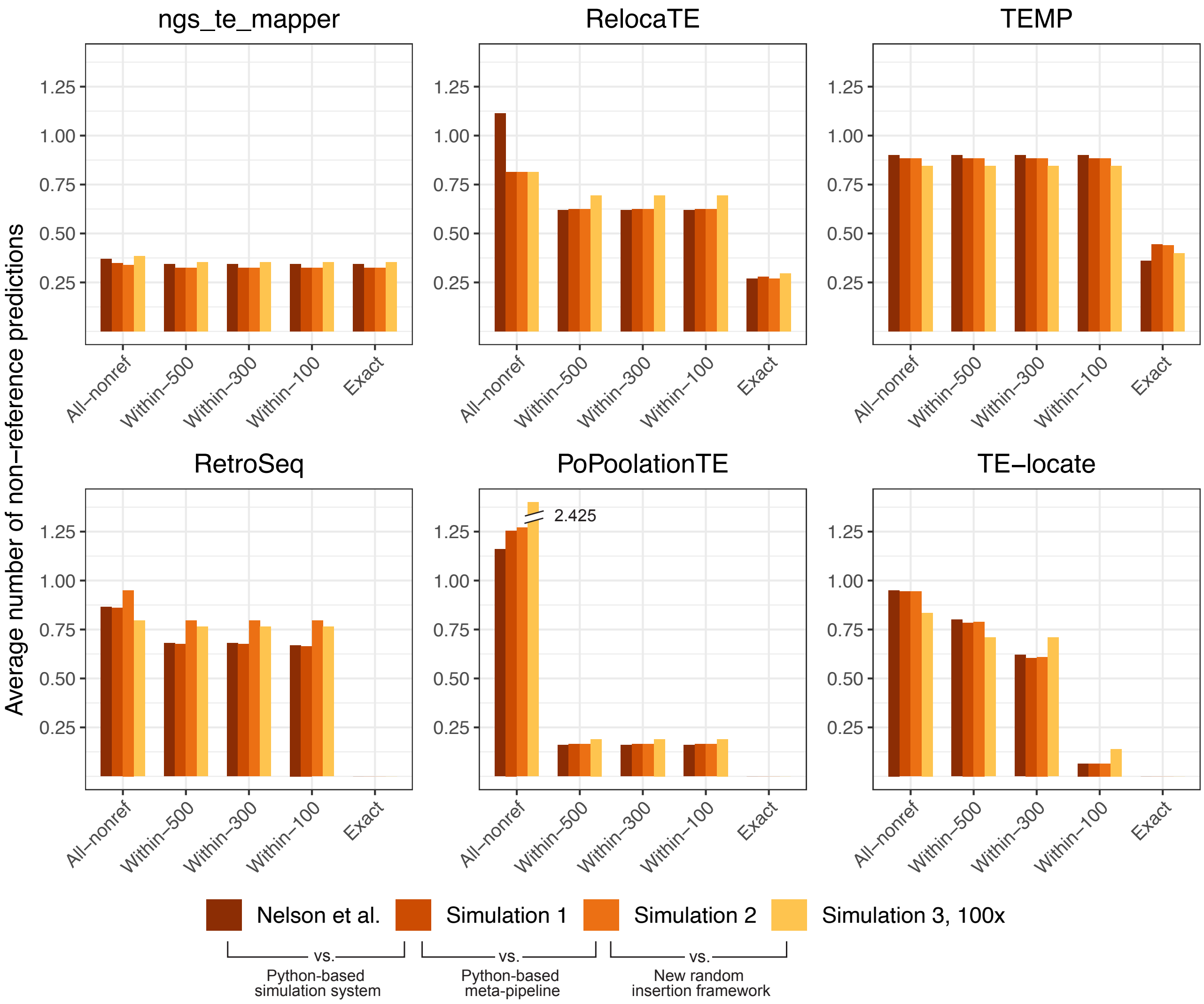


Figure S1: **Validation of McClintock 2 meta-pipeline and simulation system.** The bar plot shows the average number of all non-reference predictions and correct predictions falling in different window sizes (exact, within 100 bp, within 300 bp and within 500 bp) across all simulated replicates, including old McClintock data [1], simulation 1, simulation 2 and simulation 3 (100×). Values shown in this figure are the average number of the forward and reverse strand since two strands were run separately in our simulation framework. Only six original methods included in McClintock are shown here for validation. The PoPoolationTE value exceeds y-axis limitation and thus is annotated with a break and the actual number.

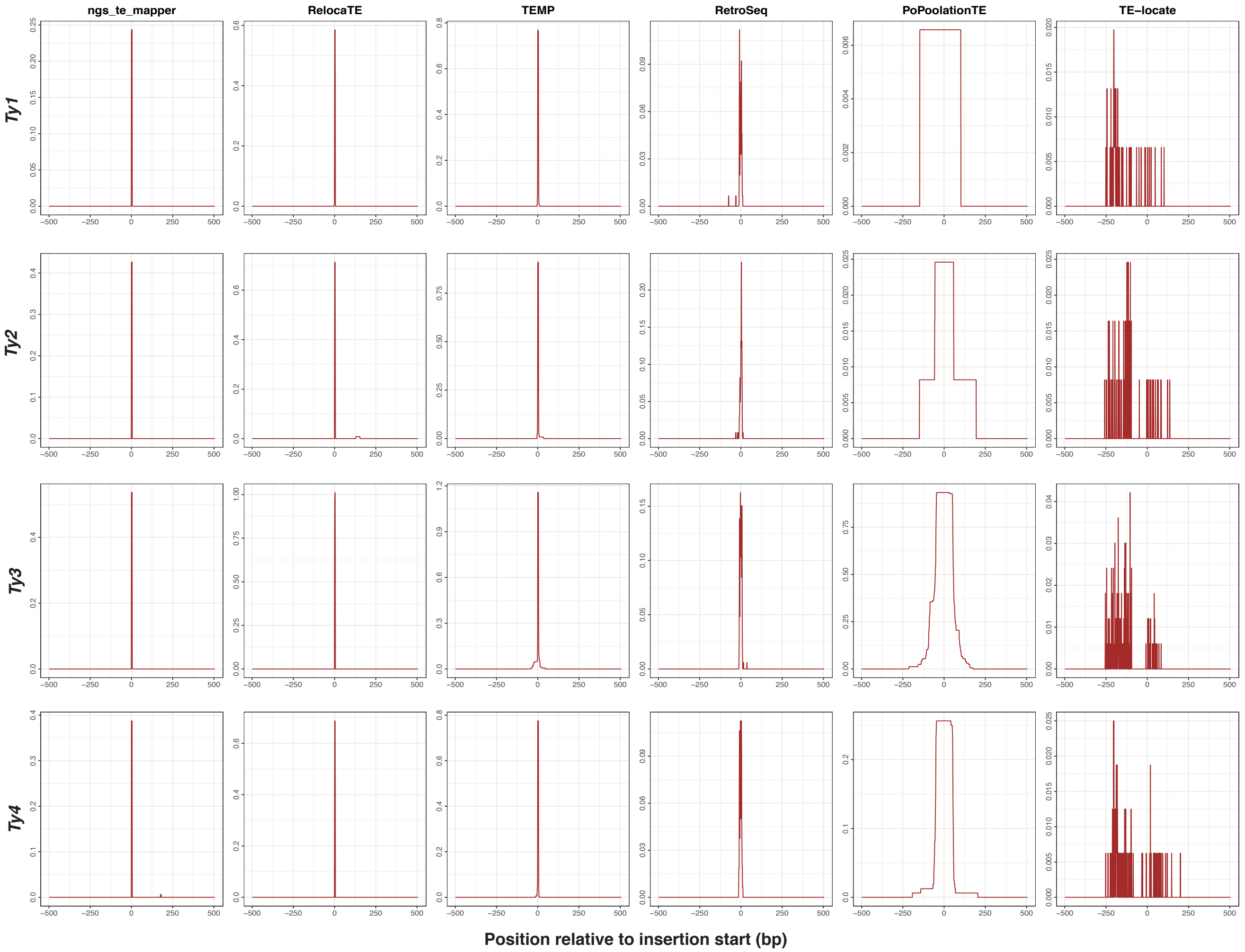


Figure S2: **Positional accuracy of six original component methods revealed by single insertion simulations that use a model of TE insertion based on yeast targeting preferences.** Plots show average coverage for non-reference predictions relative to the coordinate of the start position of synthetic insertions. This figure is based on component method predictions using 50× simulated WGS data as input. The maximum window size inspected here is 500 bp from the synthetic insertion.

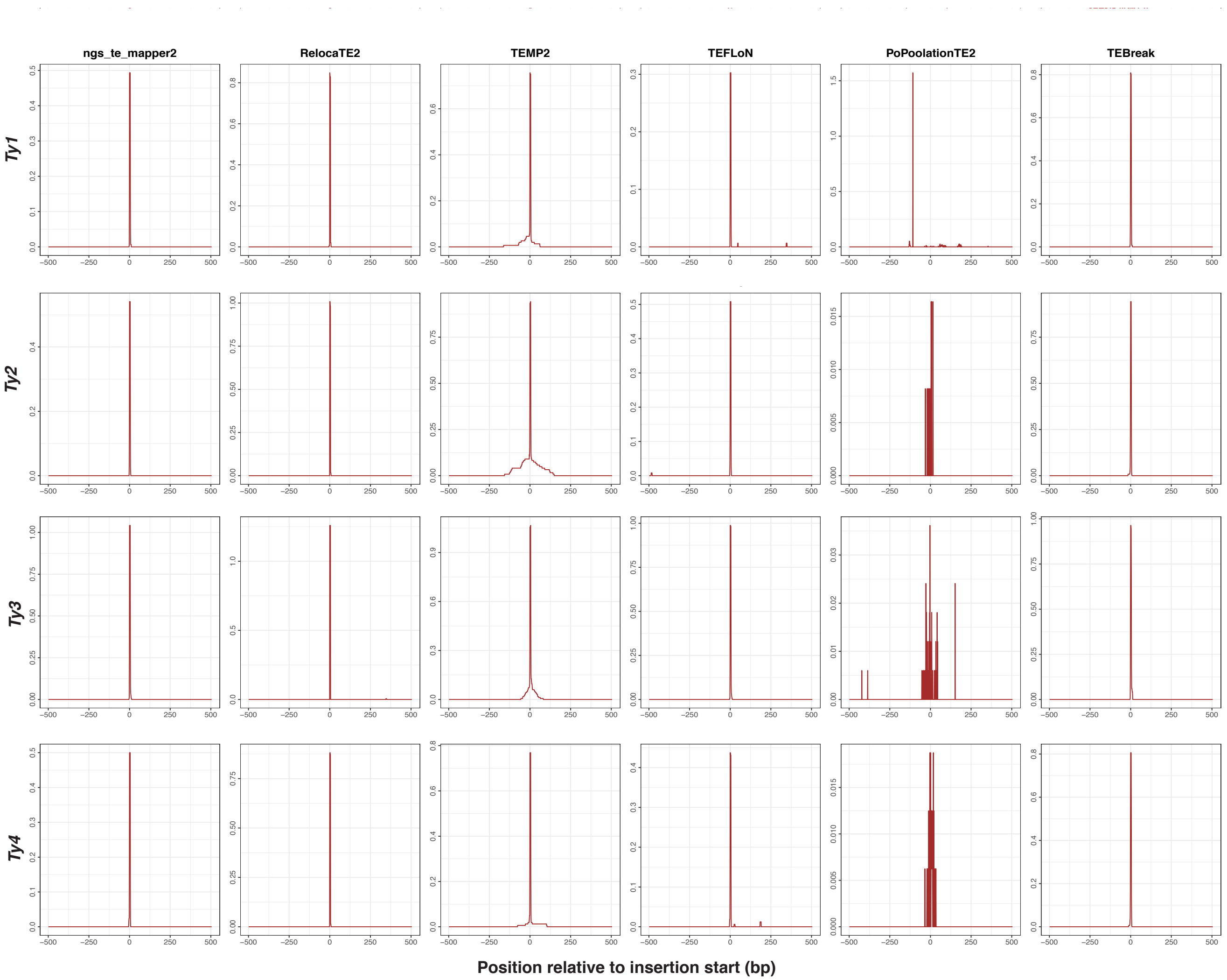


Figure S3: **Positional accuracy of six new component methods revealed by single insertion simulations that use a model of TE insertion based on yeast targeting preferences.** Plotting criteria are the same as in Figure S2.

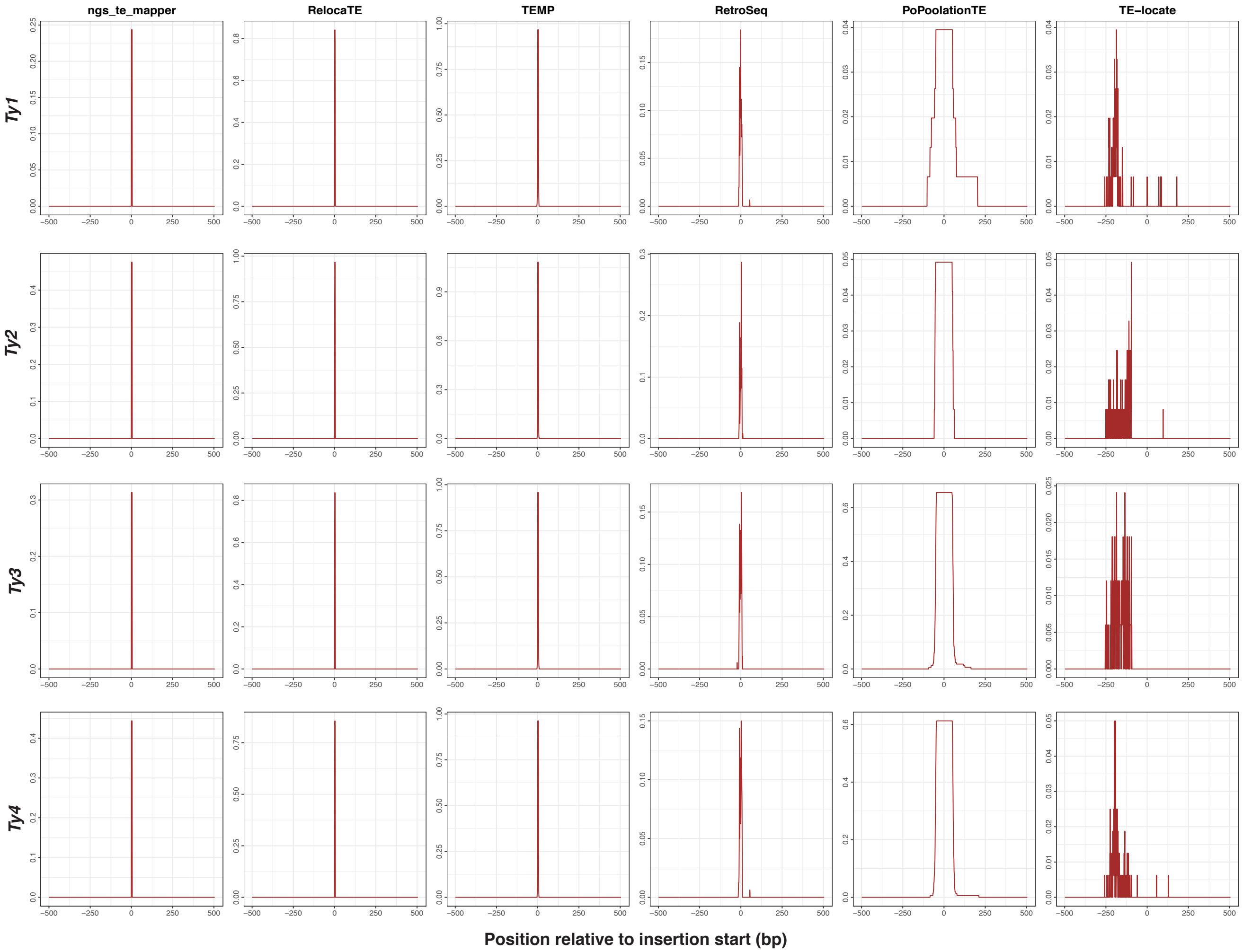


Figure S4: **Positional accuracy of six original component methods revealed by single insertion simulations that use a random model of TE insertion in non-repetitive regions of the yeast genome.** Plotting criteria are the same as in Figure S2.

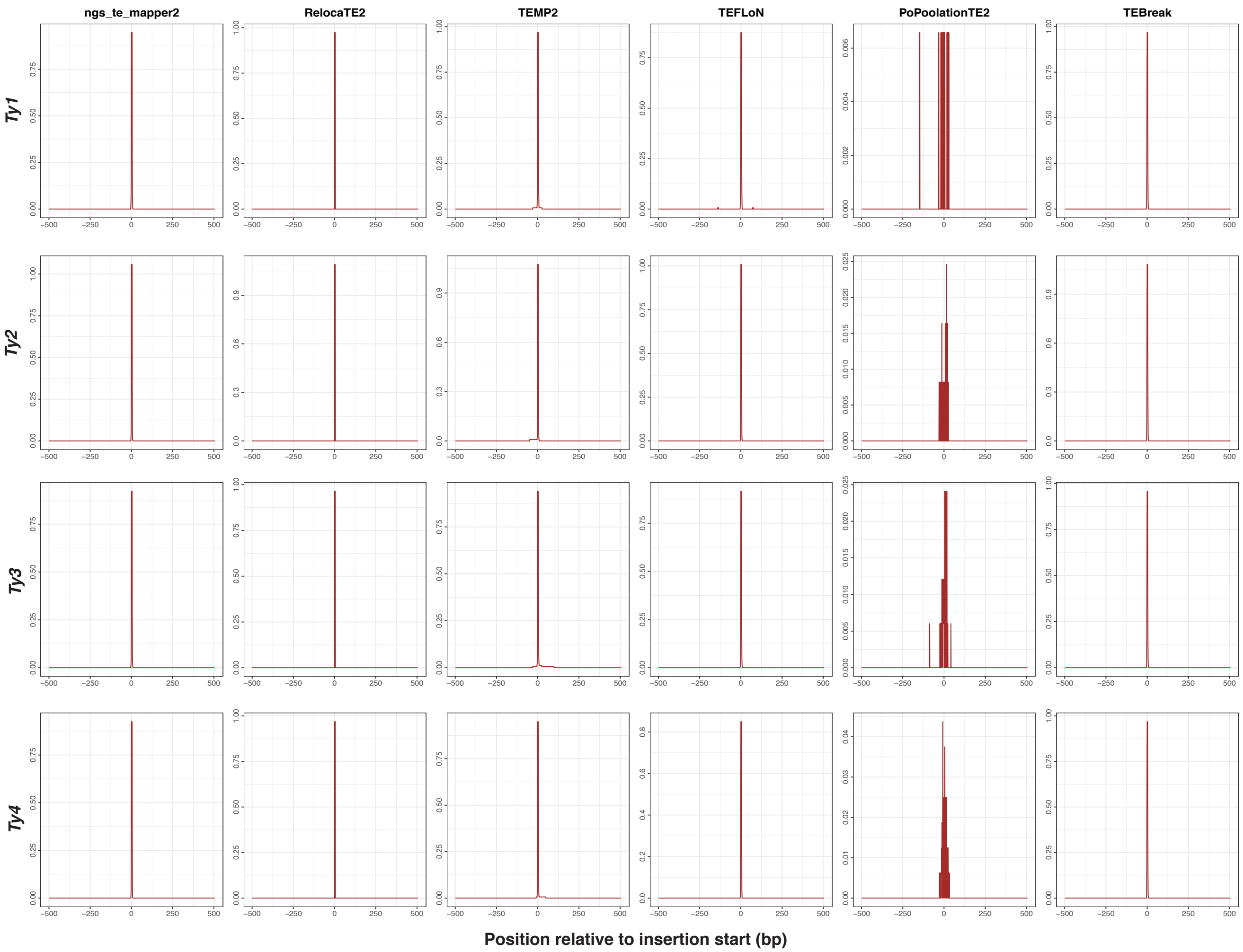


Figure S5: **Positional accuracy of six new component methods revealed by single insertion simulations that use a random model of TE insertion in non-repetitive regions of the yeast genome.** Plotting criteria are the same as in Figure S2.

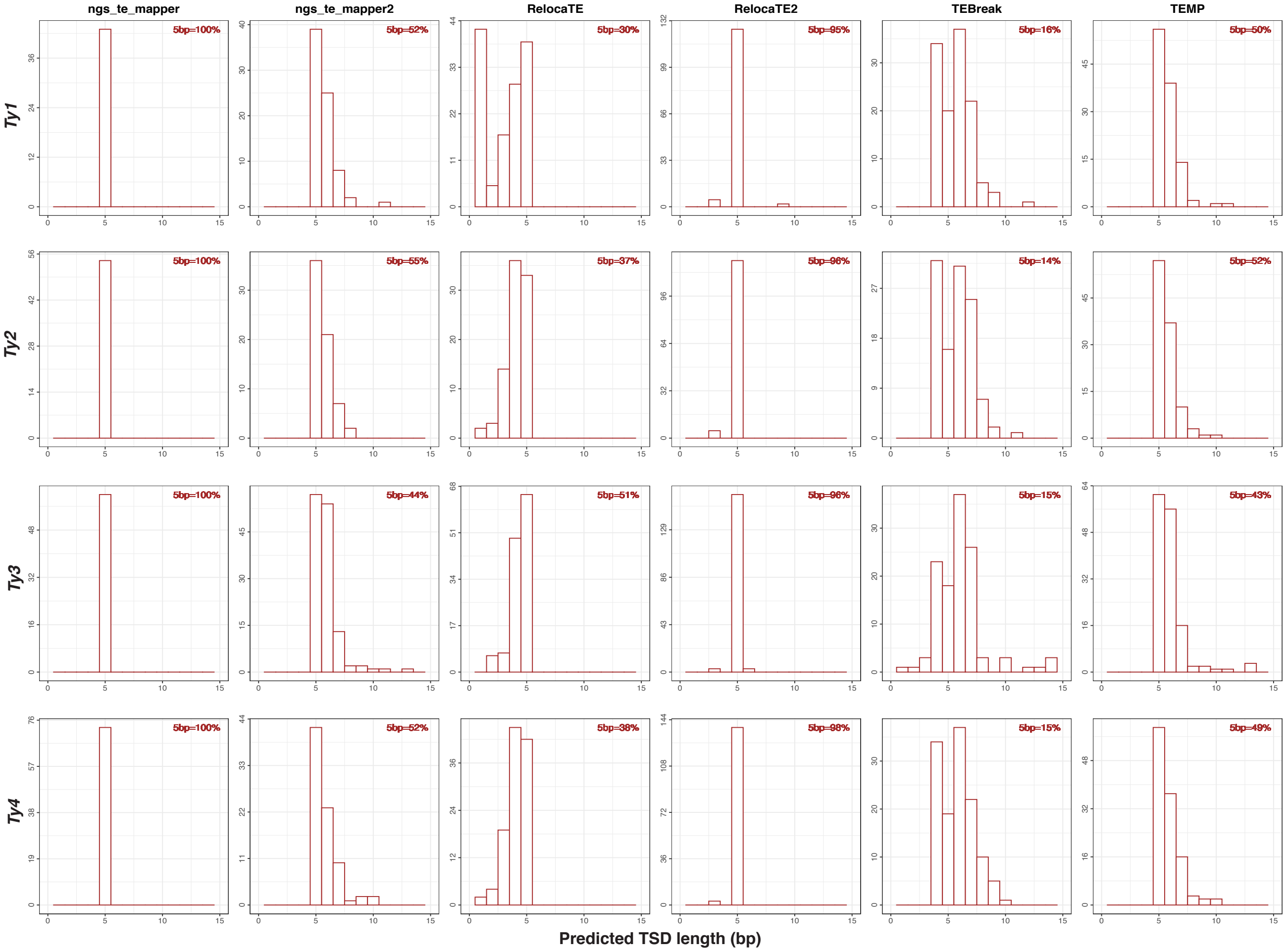


Figure S6: **TSD length distribution predicted by split-read methods in 50× yeast model simulation.** Histograms of TSD lengths predicted by split-read methods (ngs_te_mapper, ngs_te_mapper2, RelocaTE, RelocaTE2, TEBreak and partial TEMP) are plotted based on simulation 3 applying yeast model.

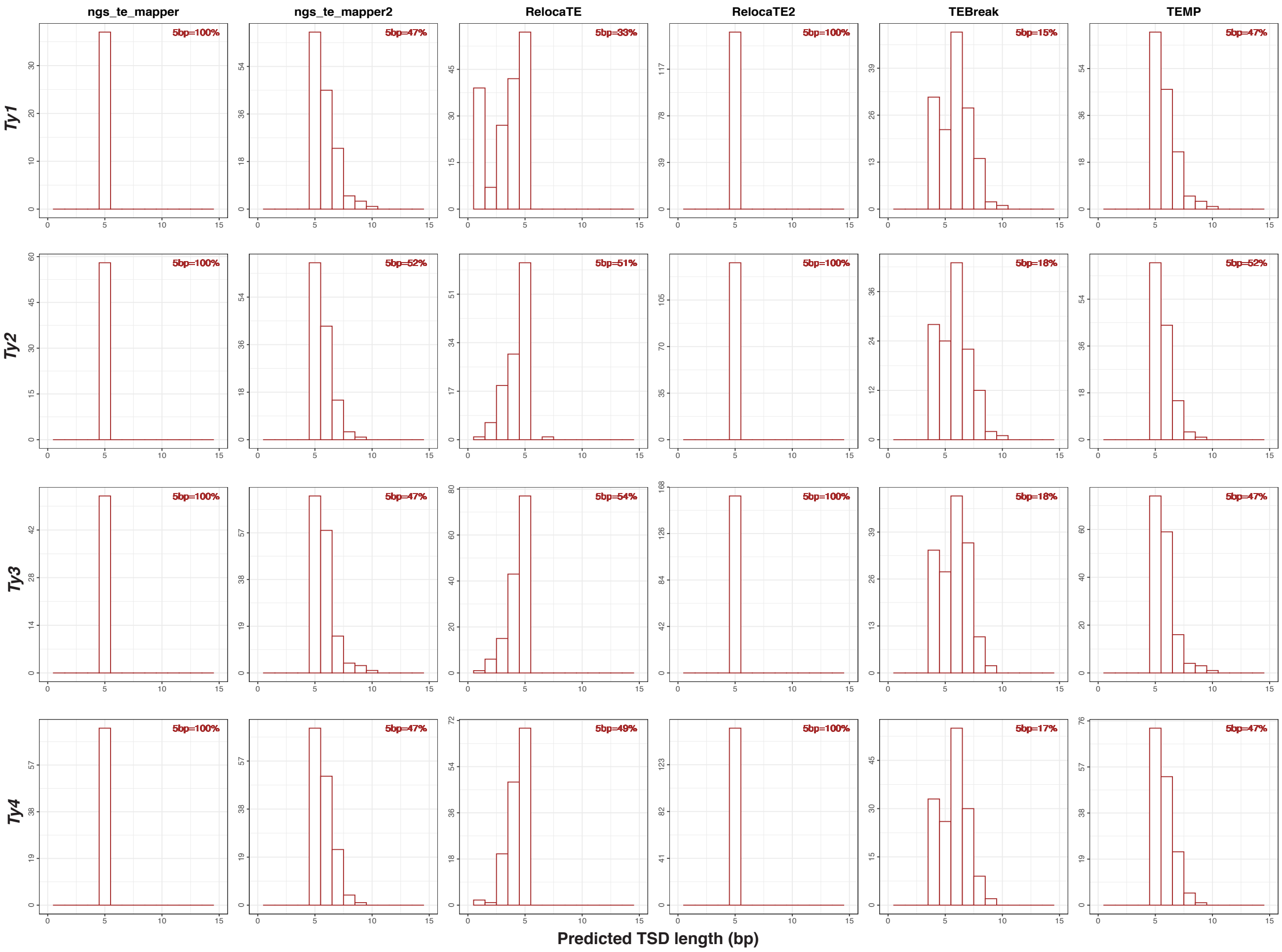


Figure S7: **TSD length distribution predicted by split-read methods in 50× random simulation.** Same as Fig S6 based on simulation 4 applying random model.

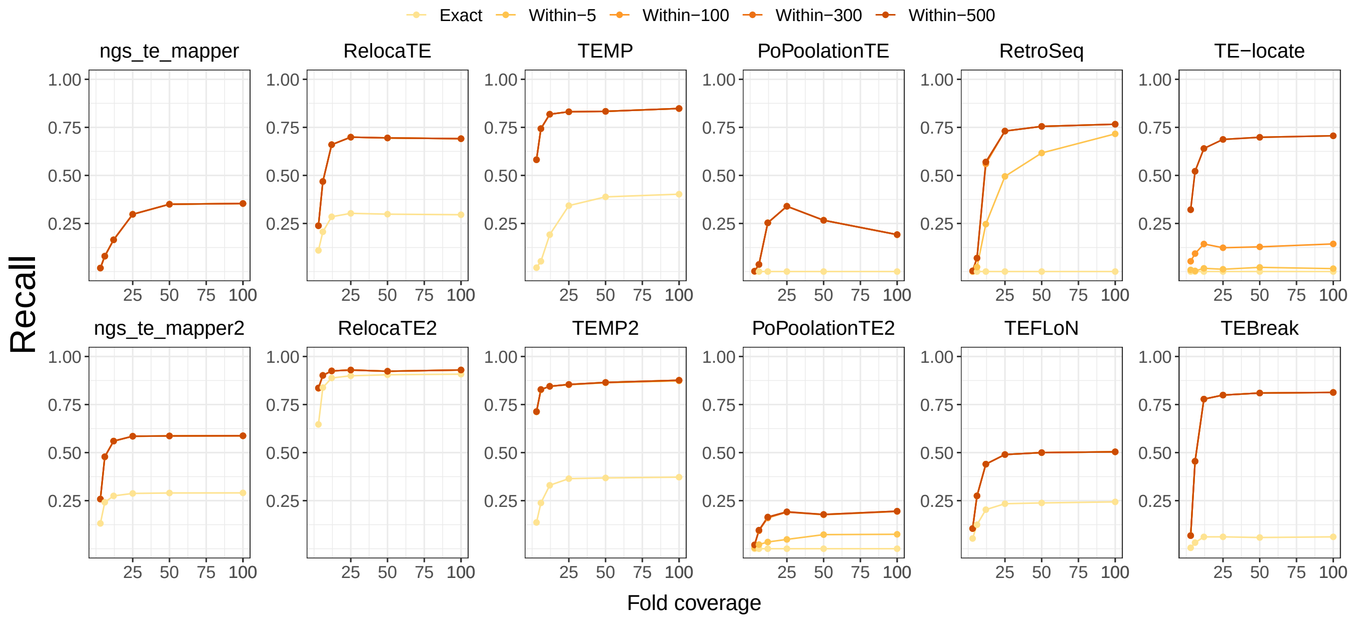


Figure S8: **Recall of 12 component methods within multiple overlapping windows in simulation of yeast model.** Shown are the recall of 12 component methods across variant synthetic fold coverage (i.e., 3×, 6×, 12×, 25×, 50× and 100×) in the simulation applying yeast model. Curves in different colors indicate variant overlapping window sizes to identify true-positive predictions, including exact coordinates, 5 bp, 100 bp, 300 bp, 500 bp. The six old component methods in McClintock are organized in the first line of each panel, while six additional methods are arranged in the second line.

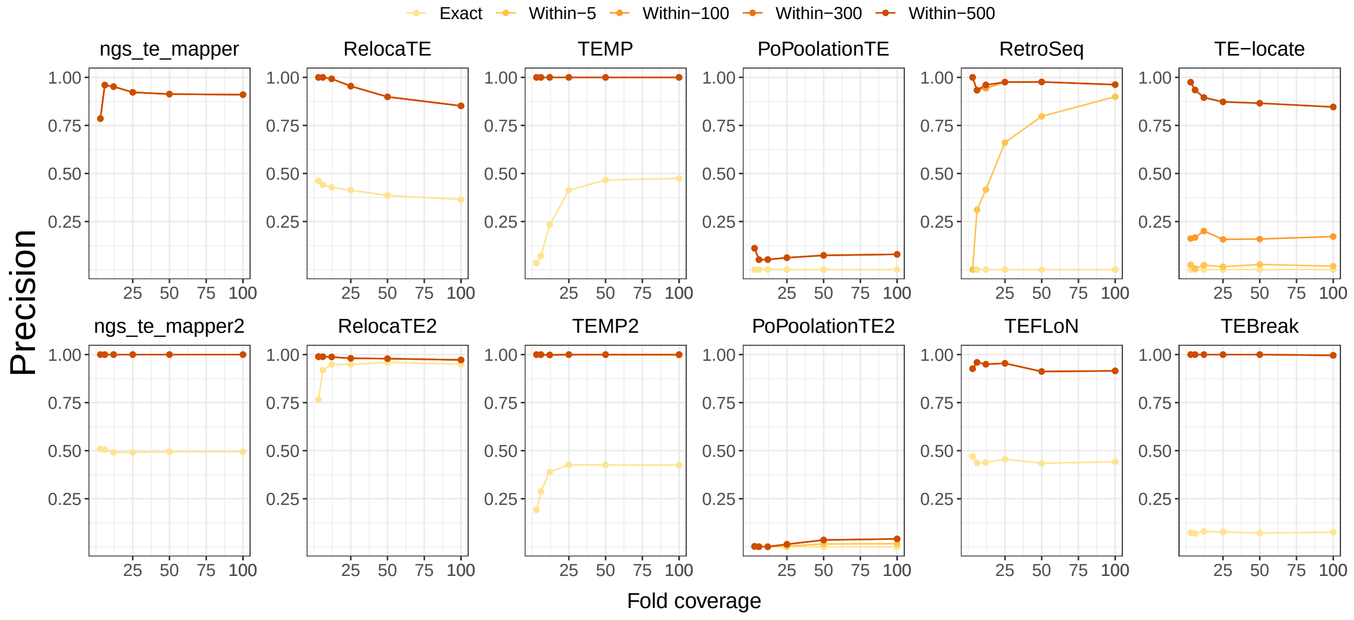


Figure S9: **Precision of 12 component methods within multiple overlapping windows in simulation of yeast model.** Precision curves with same layout as figure S8.

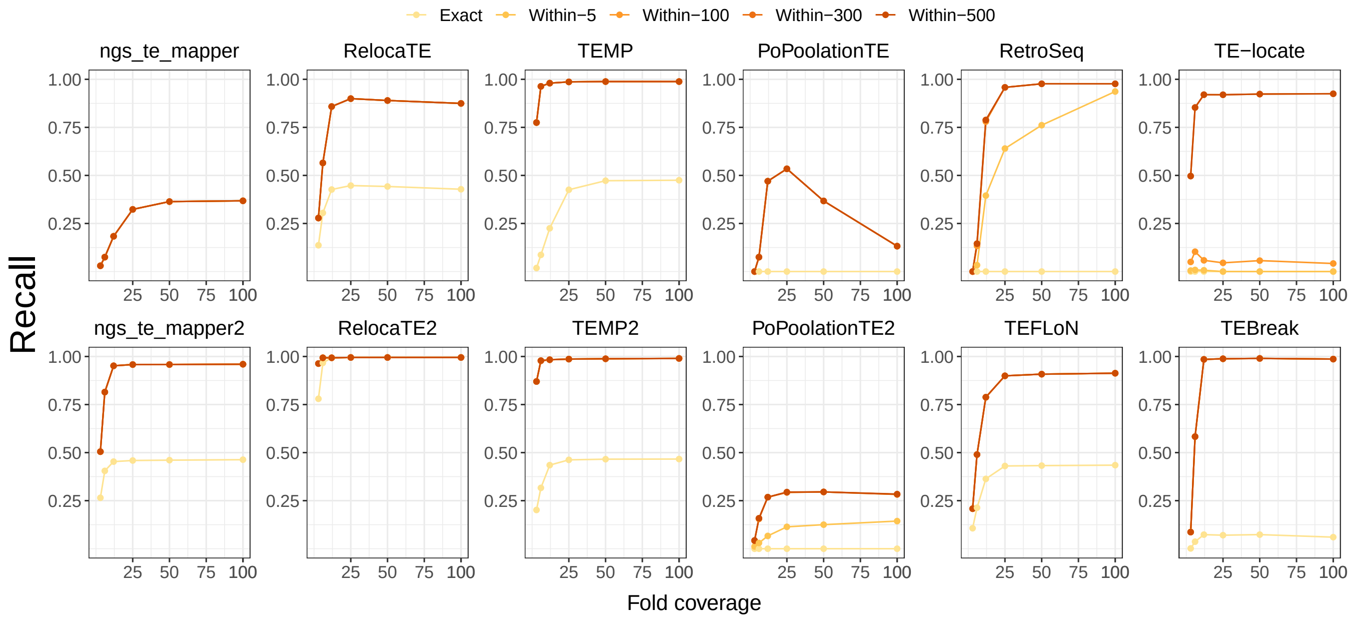


Figure S10: **Recall of 12 component methods within multiple overlapping windows in simulation of random insertion model.** Shown are recall curves based on random simulation results with same layout as figure S8.

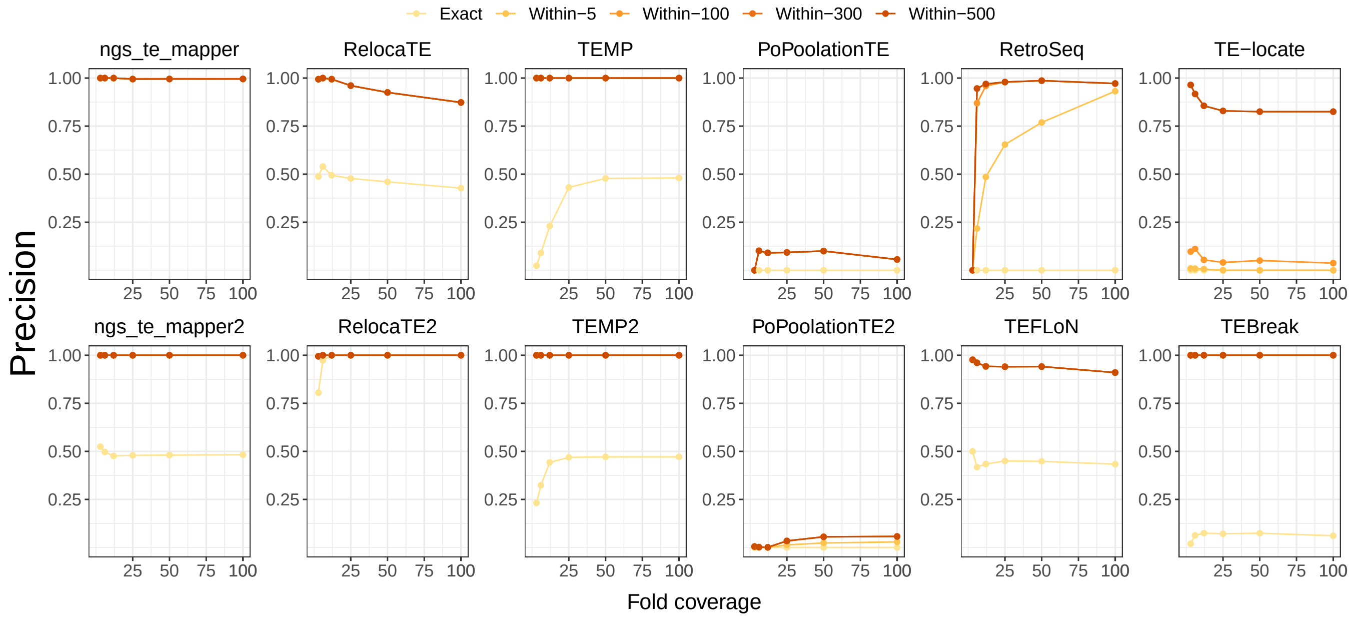


Figure S11: **Precision of 12 component methods within multiple overlapping windows in simulation of yeast model.** Shown are precision curves based on random simulation results with same layout as figure S8.

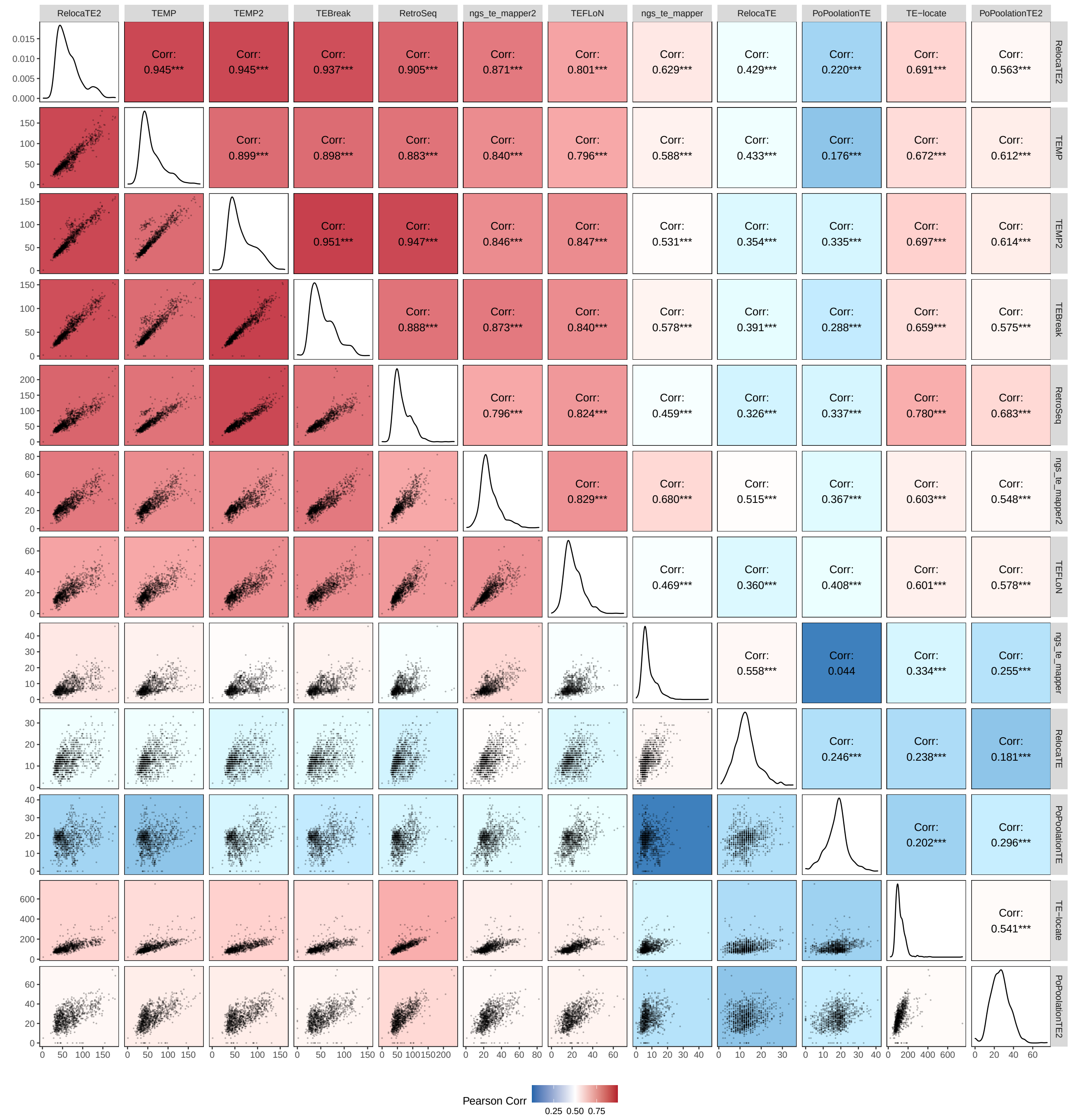


Figure S12: **Correlation across 12 component methods in all non-reference predictions for 1,011 yeast isolates.** The number of non-reference predictions for each yeast sample was plotted and compared across 12 component methods. WGS data are from [15, 16] and down-sampled to 50× fold-coverage before analysis. The panels in lower-left triangle are scatter plots on total TE count of each isolate between the corresponding pairs of methods. Panels in the upper-right triangle reveal the Pearson correlation between pairs of methods. Background colors for each panel are selected related to the Pearson correlation statistic, where higher and lower correlations are plotted in red and blue respectively. Diagonal panels show the distribution of TE counts made by each component method. The component methods are re-ordered according to the general performance revealed in our simulation results.

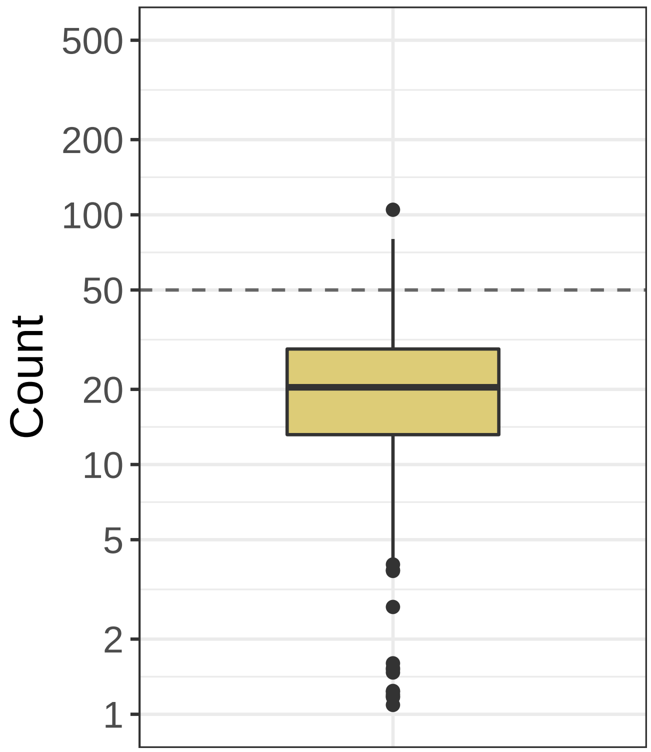


Figure S13: **Distribution of estimated copy number of full-length TEs across 1,011 yeast isolates.** Shown is the distribution of copy number of full-length TEs per strain (summed over all Ty families) estimated by the McClintock 2 coverage module using internal coding regions of all Ty families. Estimated full-length TE copy number includes both reference and non-reference TEs, but does not include solo LTRs. All WGS samples from [15, 16] were down-sampled to 50× fold-coverage prior to analysis. The y-axis is transferred to a $\log_{10}$ scale. The line inside the box indicates the median value, the colored box shows the interquartile range (IQR), whiskers show values 1.5×IQR of the upper or lower quartiles, and the dots indicate outliers that beyond 1.5×IQR.

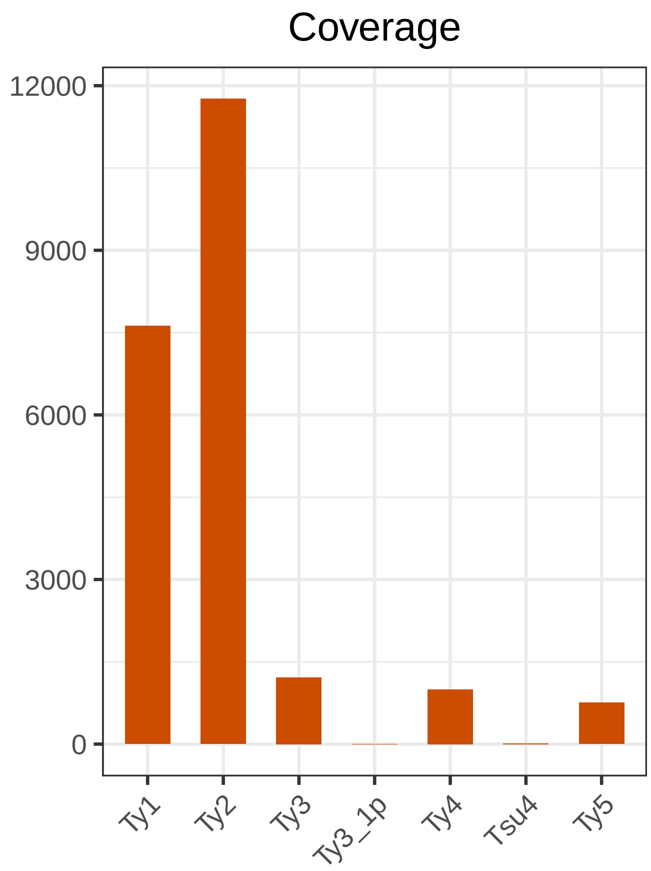


Figure S14: **Estimated total number of full-length TEs per Ty family across all 1,011 yeast isolates.** Shown is the total copy number of full-length TEs (summed across all strains) estimated by the McClintock 2 coverage module using internal coding regions of all Ty families. Estimated full-length TE copy number includes both reference and non-reference TEs, but does not include solo LTRs. All WGS samples from [15, 16] were down-sampled to 50× fold-coverage prior to analysis.

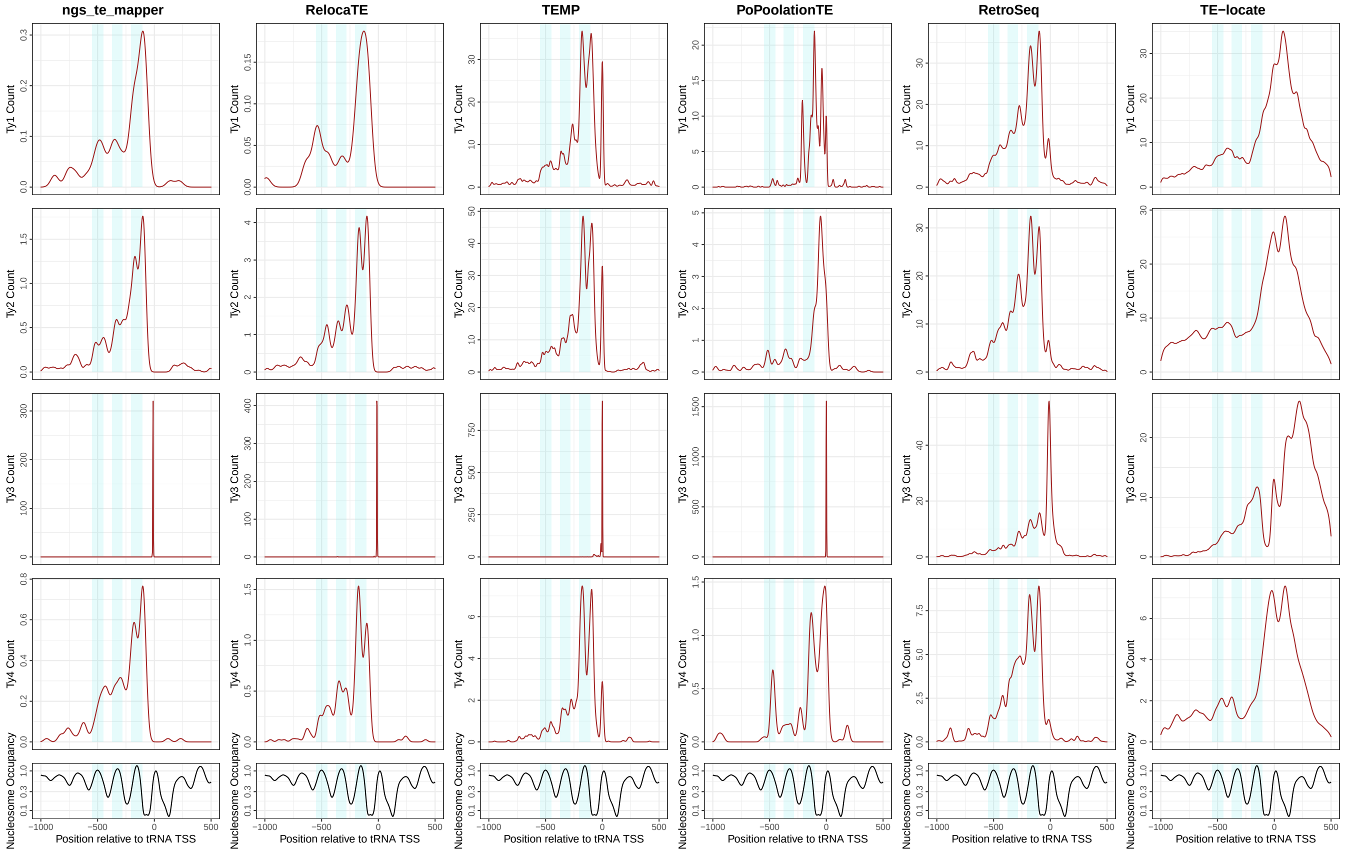


Figure S15: **TE prediction distribution relative to tRNA genes by original six methods from McClintock.** The first four rows show non-reference predictions made by six original methods in McClintock [1] from 1000 bp upstream and 500 bp downstream of tRNA genes partitioned by Ty family (summed over all strains) in 1,011 *S. cerevisiae* WGS samples [15, 16], down-sampled to 50× fold-coverage. Only four active Ty families (Fig. 4B) are included in this analysis. The bottom row shows nucleosome occupancy inferred using MNase-seq data from [17]. Light blue shades indicate 100-bp regions surrounding peaks of nucleosome occupancy.

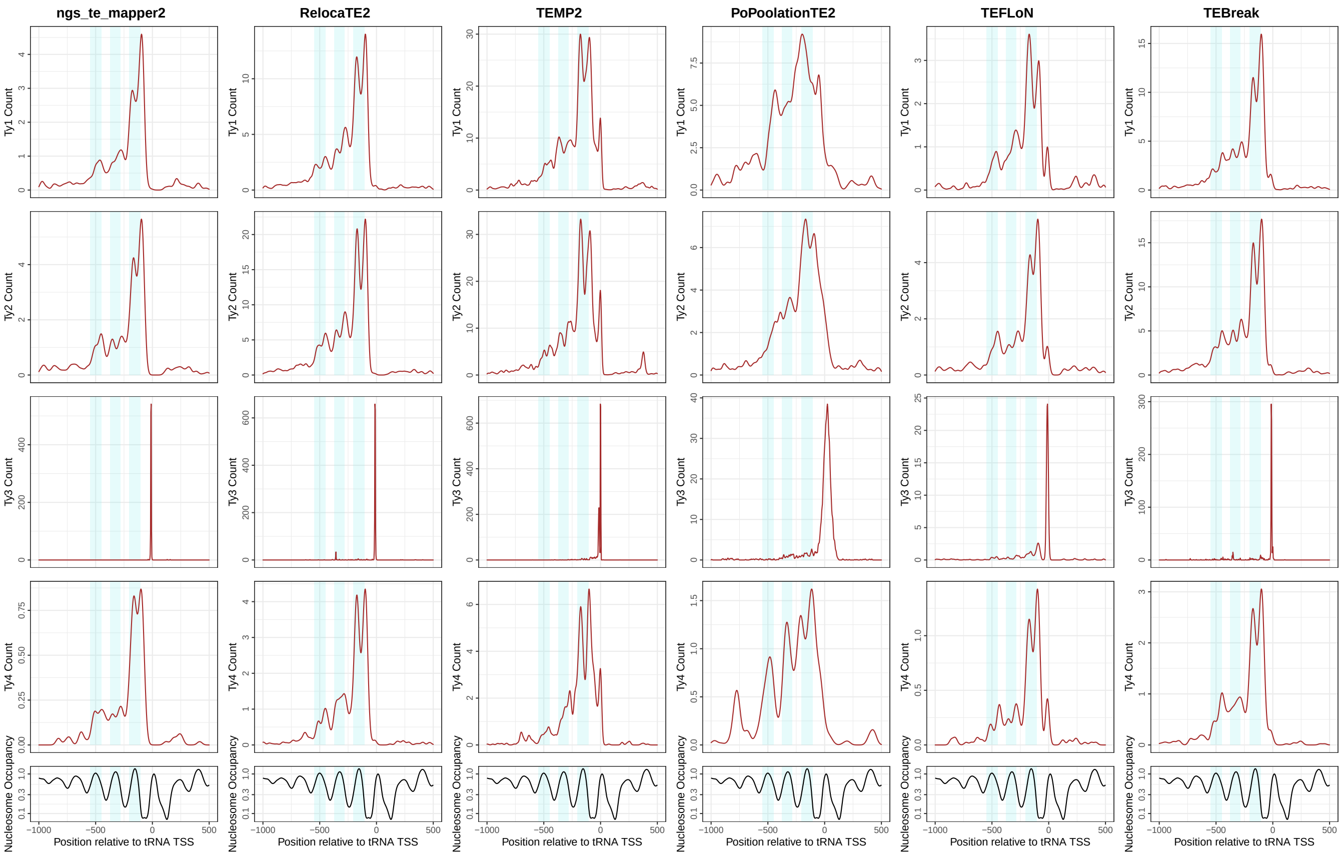


Figure S16: **TE prediction distribution relative to tRNA genes by six new methods in McClintock 2.** The first four rows show non-reference predictions made by six newly added methods in McClintock 2 from 1000 bp upstream to 500 bp downstream of tRNA genes partitioned by Ty family (summed over all strains) in 1,011 *S. cerevisiae* WGS samples [15, 16], down-sampled to 50× fold-coverage. Only four active Ty families (Fig. 4B) are included in this analysis. The bottom row shows nucleosome occupancy inferred using MNase-seq data from [17]. Light blue shades indicate 100-bp regions surrounding peaks of nucleosome occupancy.
